# pYtags enable spatiotemporal measurements of receptor tyrosine kinase signaling in living cells

**DOI:** 10.1101/2022.08.13.503850

**Authors:** Payam E. Farahani, Xiaoyu Yang, Emily V. Mesev, Kaylan A. Fomby, Caleb J. Bashor, Celeste M. Nelson, Jared E. Toettcher

**Affiliations:** Department of Chemical & Biological Engineering, Princeton University, Princeton, NJ 08544, USA; Department of Bioengineering, Rice University, Houston, TX 77030, USA; Program in Systems, Synthetic, and Physical Biology, Rice University, Houston, TX 77030, USA; Department of Molecular Biology, Princeton University, Princeton, NJ 08544, USA; Department of Biosciences, Rice University, Houston, TX 77030, USA

## Abstract

Receptor tyrosine kinases (RTKs) are major signaling hubs in metazoans, playing crucial roles in cell proliferation, migration, and differentiation. However, few tools are available to measure the activity of a specific RTK in individual living cells. Here, we present pYtags, a modular approach for monitoring the activity of a user-defined RTK by live-cell microscopy. pYtags consist of an RTK modified with a tyrosine activation motif that, when phosphorylated, recruits a fluorescently labeled tandem SH2 domain with high specificity. We show that pYtags enable the monitoring of a specific RTK on seconds-to-minutes time scales and across subcellular and multicellular length scales. Using a pYtag biosensor for epidermal growth factor receptor (EGFR), we quantitively characterize how signaling dynamics vary with the identity and dose of activating ligand. We show that orthogonal pYtags can be used to monitor the dynamics of EGFR and ErbB2 activity in the same cell, revealing distinct phases of activation for each RTK. The specificity and modularity of pYtags opens the door to robust biosensors of multiple tyrosine kinases and may enable engineering of synthetic receptors with orthogonal response programs.

**Highlights:**

- pYtags report on signaling of user-defined RTKs in living cells
- EGFR signaling dynamics depend on ligand identity and dimer strength
- rthogonal pYtags enable reporter multiplexing
- pYtags can report on signaling of endogenously expressed RTKs

## Introduction

Development and homeostasis of multicellular organisms require that cells sense and respond to diverse microenvironmental signals. Receptor tyrosine kinases (RTKs) are one widely-expressed class of cell-surface receptors that play a key role in this information processing (Lemmon and Schlessinger, 2010). RTK activation is triggered by growth factors or hormones that bind to the extracellular domains of receptors, inducing conformational changes that lead to receptor dimerization and the autophosphorylation of tyrosine residues in the C-terminal tails. Phosphorylated RTKs, hereafter described as “activated” RTKs, present phosphotyrosine-containing motifs that bind to downstream effectors that signal to multiple pathways. RTKs are thus positioned as the uppermost node of a complex network of intracellular signaling.

RTK signaling provides a high-dimensional input space through which this class of receptors regulates a variety cellular processes (Lemmon and Schlessinger, 2010). Human cells express at least 58 RTKs whose binding partners vary widely, so the pathways that are activated in a particular cell depend on the set of receptors expressed on its surface (Madhani, 2001; Salokas et al., 2022). Some classes of receptors can form homo- or heterodimers, further diversifying signaling responses (Del Piccolo et al., 2017; Harari and Yarden, 2000; Kanakaraj et al., 1991). Moreover, a single RTK can bind to many different ligands that are capable of inducing distinct conformational changes to tune downstream signaling dynamics (Freed et al., 2017; Hu et al., 2022). These observations underscore the substantial complexity present at even the uppermost node of the RTK signaling network: the interaction between receptors and their ligands.

Recent advances in live-cell biosensors have enabled the study of intracellular signaling at unprecedented spatiotemporal resolution. Signaling responses can be measured with a temporal precision on the order of seconds, and recent studies have described approaches to multiplex biosensors of multiple pathways in a single cell (Regot et al., 2014) or to use barcoding strategies to monitor many biosensors within a population of cells (Kaufman et al., 2022; Yang et al., 2021). Yet the development of biosensors for specific RTKs has been somewhat limited. A fluorescent Grb2-based biosensor has been widely used to measure general RTK activity but does not distinguish between the many receptors that can recruit Grb2 (Reynolds et al., 2003).

FRET-based biosensors of epidermal growth factor receptor (EGFR) (Itoh et al., 2005; Komatsu et al., 2011; Kurokawa et al., 2001; Sorkin et al., 2000) and platelet-derived growth factor receptor (PDGFR) (Seong et al., 2017) have been described, but rely on components of endogenous substrates that participate in interactions with multiple RTKs, and their specificity remains to be characterized. A more specific Src homology 2 (SH2)-based biosensor for EGFR was recently reported but required extensive engineering and may compete with other SH2- containing proteins for binding to phosphotyrosine motifs required for signaling (Tiruthani et al., 2019). Consequently, the field still lacks modular live-cell biosensors to monitor the activity of any specific RTK of interest.

Here, we describe pYtags, a versatile and modular RTK biosensing strategy. In the pYtag approach, the C-terminus of a user-defined RTK is labeled with a tyrosine activation motif that, when phosphorylated, binds selectively to a fluorescently labeled tandem SH2 (tSH2) reporter.

This binding results in the depletion of the fluorescent reporter from the cytosol and local accumulation at membranes where the receptor is activated. We show that an EGFR pYtag biosensor quantitatively reports on the activity of EGFR with undetectable crosstalk from other unlabeled RTKs. We use pYtags to quantify EGFR activation in response to various ligands, revealing ligand- and dose-specific signaling dynamics. Mathematical modeling and experimental validation reveal that different ligands affect the signaling dynamics of EGFR by altering the dimerization affinity of ligand-bound receptors. We further demonstrate that pYtags can be applied to other receptors including FGFR1 and the ligandless RTK ErbB2, a challenging case study due to its need to signal through receptor heterodimers. We describe a second, orthogonal pYtag that can be used to simultaneously monitor EGFR and ErbB2 activity within the same cell. Finally, we demonstrate that pYtags can be inserted into the genome using CRISPR/Cas9-based editing, which enables reporting of endogenous receptors and eliminates confounding effects from ectopic expression. pYtags thus offer a modular strategy for measuring the activity of specific RTKs in individual living cells and may serve as a platform for engineering phosphotyrosine-based cargo that can be recruited to synthetic receptors.

## Results

### An orthogonal tag for monitoring tyrosine phosphorylation events

Many biosensors of kinase activity are based on two components: a synthetic substrate that is predominantly phosphorylated by a single kinase of interest, and a domain that binds specifically to the phosphorylated substrate. To apply similar logic in the design of an RTK biosensor, it would be desirable to introduce a tyrosine-containing peptide that can be uniquely phosphorylated by an RTK of interest and a binding domain that interacts only with the phosphorylated peptide. Fulfilling these requirements is especially challenging in the case of tyrosine kinases, which typically lack one-to-one specificities between tyrosine kinase, tyrosine substrate, and SH2-binding domains (Miller, 2003). However, immune cells offer examples of highly specific phosphotyrosine signaling. During activation of the T-cell receptor (TCR), pairs of tyrosine residues located in immunoreceptor tyrosinase-based activation motifs (ITAMs) are phosphorylated and serve as sites for the selective recruitment of the tandem SH2 domain of ZAP70 (ZtSH2), which binds 100-fold more tightly than individual ZAP70 SH2 domains with the same ITAMs and displays minimal crosstalk with other phosphotyrosine-containing peptides (**Figure 1A**) (Isakov et al., 1995; Labadia et al., 1996; Wange et al., 1993). Because neither the TCR nor ZAP70 are expressed by non-immune cells, we reasoned that an ITAM/ZtSH2 pair may be repurposed as an orthogonal interaction module to monitor the activity of a desired RTK in non-immune contexts. If ITAMs appended to the C-terminal tail of an RTK are phosphorylated upon activation of the RTK, this should create a selective binding site for a fluorescently labeled ZtSH2 (**Figure 1B**). At the cellular level, such a ZtSH2-based reporter should reside in the cytosol when RTK signaling is low and be recruited to the cell membrane when RTK signaling is high (**Figure 1B**).

**Figure 1.**
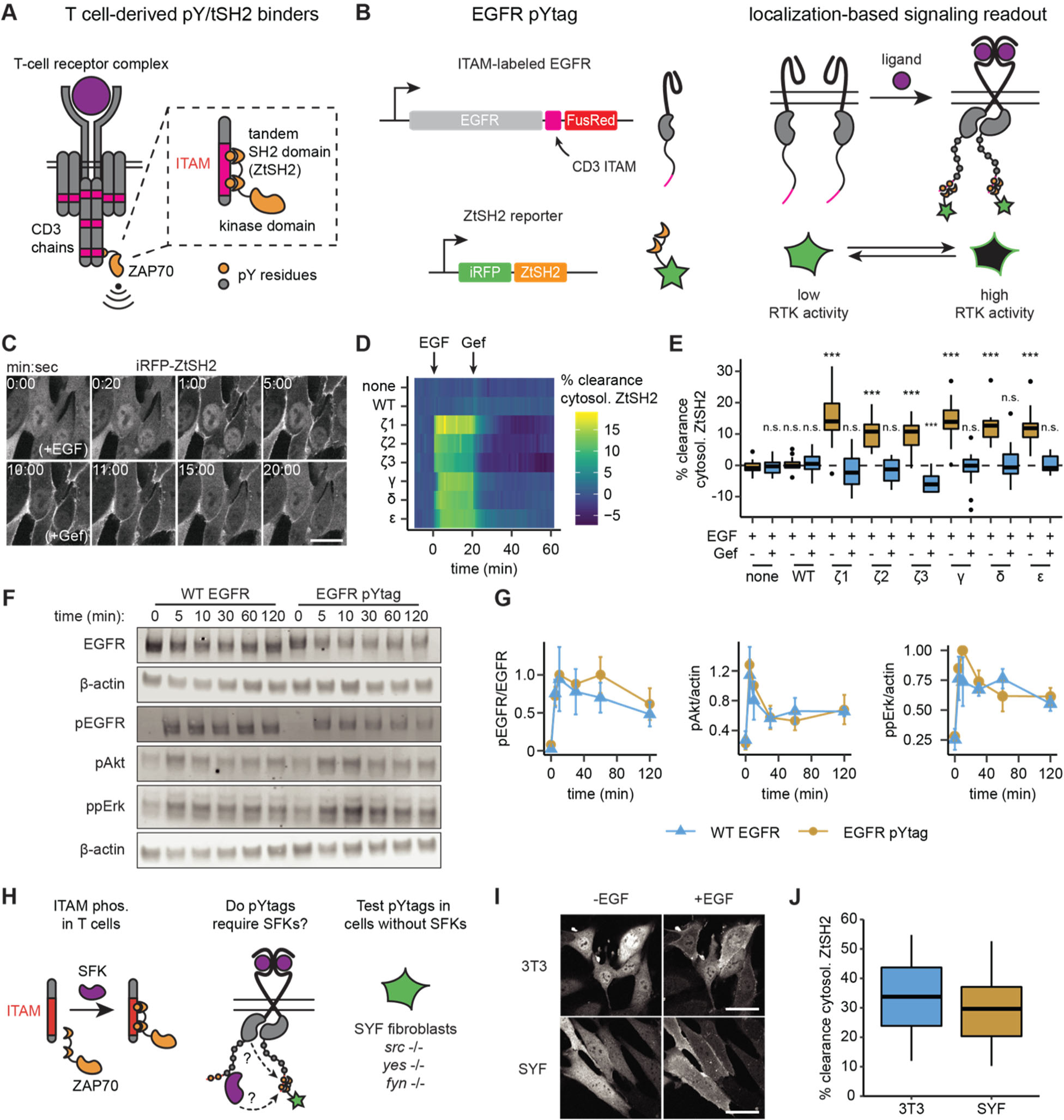
pYtags: a biosensing strategy to monitor RTK activity in living cells. (**A**) The T- cell receptor complex contains six ITAMs from CD3 chains that, when phosphorylated, bind to the tSH2 domain of ZAP70 (ZtSH2). (**B**) Three repeats of CD3 ITAMs were appended to the C- terminus of EGFR and clearance of ZtSH2 from the cytosol was assessed. (**C**) Timelapse images of NIH 3T3 cells expressing EGFR pYtag (CD3ε variant), treated first with EGF (100 ng/mL) and then with Gefitinib (10 µM). Scale bar, 20 µm. (**D**) Mean clearance of cytosolic ZtSH2 in cells co-expressing iRFP-ZtSH2 and EGFR C-terminally labeled with one of six CD3 ITAMs. EGF (100 ng/mL) and Gefitinib (10 µM) were sequentially added at times denoted by arrows. *n* = 2 independent experiments. (**E**) Clearance of cytosolic ZtSH2 10 min post-EGF treatment and 40 min post-Gefitinib treatment from (D). Lines denote mean values, boxes denote 25-75^th^ percentiles, and whiskers denote minima and maxima. *n ≥* 14 cells from 2 independent experiments. n.s. not significant, *** *p* < 0.001 by Kolmogorov-Smirnov test with cells expressing no additional EGFR 10 min post-EGF. (**F**) Immunoblots of NIH 3T3 cells expressing either WT EGFR or EGFR pYtag treated with EGF (100 ng/mL). (**G**) Mean ± S.E.M. levels of EGFR, Akt, and Erk phosphorylation from (F). *n* = 3 independent experiments. (**H**) The EGFR pYtag was tested in SYF cells to determine whether SFKs are required for ITAM phosphorylation. (**I**) Representative images of NIH 3T3 and SYF cells expressing EGFR pYtag, treated with EGF (100 ng/mL). Scale bars, 40 µm. (**J**) Mean clearance of cytosolic ZtSH2 in SYF and NIH 3T3 cells 10 min after treatment with EGF. For each condition, *n* > 20 cells from 2 independent experiments. See also Video S1.

We first applied this biosensing approach to detect the activation of EGFR, the most well-characterized of the 58 known human RTKs (Salokas et al., 2022). We tested six ITAMs from the CD3γ, CD3δ, CD3ε, and CD3ζ chains of the TCR. In each case, we fused three identical repeats of the ITAM to the C-terminal tail of EGFR followed by the FusionRed fluorescent protein (EGFR-ITAM-FusionRed). We then co-expressed each receptor with an iRFP-labeled ZtSH2 (iRFP-ZtSH2) in NIH 3T3 fibroblasts, which express negligible levels of EGFR (**Figure 1B**) (Di Fiore et al., 1987). In cells expressing EGFR-ITAM-FusionRed, treatment with epidermal growth factor (EGF) resulted in rapid translocation of ZtSH2 from the cytosol to the cell membrane (**Figure 1C; Video S1**), which we assessed by quantifying the percentage of ZtSH2 cleared from the cytosol (**see Method details**). Clearance of ZtSH2 from the cytosol was quickly reversed by treatment with Gefitinib, an inhibitor of EGFR kinase activity, indicating that ZtSH2 can serve as a rapid, reversible reporter of EGFR activation (**Figure 1C**). Cells lacking EGFR or expressing an ITAM-less EGFR-FusionRed exhibited no change in ZtSH2 localization in response to treatment with EGF or Gefitinib (**Figures 1D and 1E**), demonstrating that the ZtSH2 reporter responds specifically to the activation of an ITAM- labeled RTK. Furthermore, a Grb2-based reporter (Reynolds et al., 2003) localized to both EGFR-CD3ε-FusionRed and EGFR-FusionRed in cells stimulated with EGF, confirming that the Grb2-based reporter cannot discriminate between ITAM-labeled and ITAM-less RTKs (**Figure S1**). Although all six ITAMs that we tested appear capable of functioning as biosensors of RTK activity, we chose to focus on the CD3ε ITAM for all subsequent experiments due to its reported selectivity for ZtSH2 over other phosphotyrosine-binding domains (Love and Hayes, 2010; Osman et al., 1995; Ravichandran et al., 1993). We refer to the resulting two-component biosensor as a phosphotyrosine tag (pYtag): a tyrosine activation motif that recruits its complementary tSH2 reporter to report on signaling.

Since pYtags introduce additional tyrosine residues and SH2-containing peptides to an RTK signaling complex, we asked whether this biosensing strategy interferes with signaling downstream of EGFR. We stimulated NIH 3T3 cells expressing either EGFR-FusionRed or EGFR pYtag (EGFR-CD3ε-FusionRed; iRFP-ZtSH2) with EGF and measured signaling responses as a function of time by immunoblotting (**Figure 1F**). We found that EGFR, Erk, and Akt were phosphorylated at similar levels in the two cell lines (**Figure 1G**), suggesting that ITAM phosphorylation and ZtSH2 recruitment do not interfere with signaling downstream of EGFR.

In T cells, CD3 ITAMs are typically phosphorylated by Src family kinases (SFKs) upon engagement of TCRs (Gaud et al., 2018). Because EGFR also activates SFKs, we reasoned that pYtags could be phosphorylated indirectly by EGFR-associated SFKs rather than by the kinase domain of EGFR itself (**Figure 1H**). To test whether SFK-mediated phosphorylation is required for pYtag function, we expressed EGFR pYtag in mouse embryonic fibroblasts lacking all three ubiquitously expressed SFKs: Src, Yes, and Fyn (SYF cells) (**Figure 1H**) (Klinghoffer et al., 1999). As in NIH 3T3 cells, SYF cells exhibited strong clearance of ZtSH2 from the cytosol after stimulation with EGF (**Figures 1I and 1J**). Combined with the rapid responses observed after EGFR stimulation and inhibition (**Figures 1C-1E**), these results suggest that the EGFR pYtag acts as a direct biosensor of EGFR activity.

### pYtags reveal the dynamics of EGFR signaling at high spatiotemporal resolution

Because ZtSH2 can in principle be recruited rapidly on and off the membrane and may localize to distinct subcellular compartments, we reasoned that pYtags could be used to monitor the subcellular activity of an RTK in individual cells over time. To test this possibility, we first performed rapid, high-resolution imaging of NIH 3T3 cells treated with EGF (**Figure 2A**). We found that ZtSH2 was cleared from the cytosol in a biphasic manner, with an initial phase of rapid cytosolic clearance occurring within the first ∼40 sec, followed by a further gradual increase in cytosolic clearance over the subsequent ∼20 min (**Figure 2B**). Similar biphasic responses for EGFR were observed previously at the population-level using a split luciferase system (Macdonald-Obermann and Pike, 2014; Macdonald-Obermann et al., 2012) and *in vitro*phosphorylation assays (Kovacs et al., 2015), but have never been reported for individual cells.

**Figure 2.**
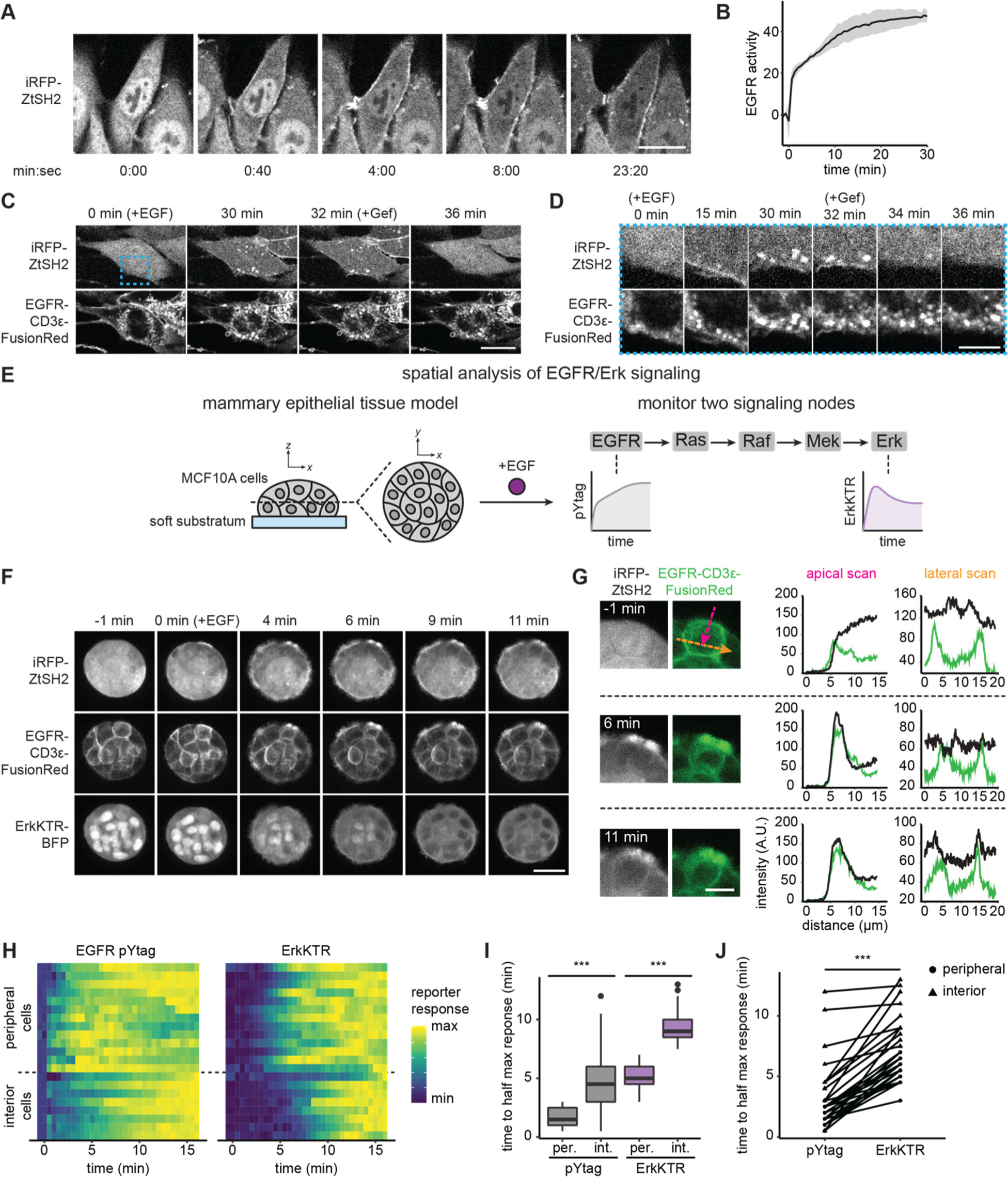
Monitoring EGFR signaling at subcellular and multicellular length scales. (**A**) Images of EGFR pYtag-expressing NIH 3T3 cells treated with EGF (20 ng/mL). Scale bar, 20 µm. (**B**) Mean ± S.D. clearance of cytosolic ZtSH2 from (A). *n* = 3 independent experiments. (**C**) EGFR pYtag-expressing NIH 3T3 cells treated with EGF (20 ng/mL) were monitored for internalized ZtSH2-positive vesicles, and then treated with Gefitinib (10 µM). Scale bar, 20 µm. (**D**) Timelapse images from the region denoted by the blue dashed border from (C). Scale bar, 10 µm. (**E**) MCF10A human mammary epithelial cells cultured on soft substrata form round, multilayered clusters. EGFR pYtag and ErkKTR were used to spatiotemporally monitor both EGFR and Erk responses after stimulation with EGF. (**F**) Images of MCF10A cells cultured on soft substrata and treated with EGF (100 ng/mL). Scale bar, 25 µm. (**G**) Apical and lateral enrichment of ZtSH2 was quantified by line scans denoted by magenta and orange vectors, respectively. Scale bar, 10 µm. (**H**) Heatmaps of EGFR pYtag and ErkKTR responses from (F). Rows denote individual cells. For each cell, signaling responses of each biosensor were normalized to their respective minima and maxima. (**I**) Time to half maximal response for EGFR pYtag and ErkKTR for cells from (H). *n* = 19 (periphery) and *n* = 11 (interior) cells from 3 biological replicates. (**J**) Time to half maximal response for EGFR pYtag and ErkKTR. Responses of individual cells are denoted by points and connected by lines. *n* = 30 cells from 3 biological replicates. *** *p* < 0.001 by Kolmogorov-Smirnov test. See also Video S2.

Over longer time scales, EGFR can be trafficked through internalization, degradation, and recycling back to the cell membrane (Sorkin and Goh, 2009). Notably, we observed that pYtag-expressing cells stimulated for at least 30 min with EGF contained internalized vesicles that were positive for both total EGFR and ZtSH2 (**Figures 2C and 2D**). Subsequent treatment with Gefitinib eliminated ZtSH2 from EGFR-positive vesicles within minutes, suggesting that the enrichment of ZtSH2 at vesicles is indicative of signaling from endosomal compartments (**Figures 2C and 2D**). These results are consistent with prior reports that internalized EGFR remains bound to its ligand and can transmit signals from endosomal compartments (Haugh et al., 1999).

At the tissue scale, the spatiotemporal distribution of RTK signaling can also be regulated by several processes including paracrine signaling (De Simone et al., 2021), morphogen gradients (Casanova and Struhl, 1989; Sprenger and Nüsslein-Volhard, 1992), and the mechanical properties of the local microenvironment (Farahani et al., 2021). We therefore asked whether pYtags could be used to monitor RTK signaling in multicellular contexts. We previously found that MCF10A human mammary epithelial cells form multicellular clusters when cultured on soft substrata and exhibit a complex spatial pattern of EGF binding at cell membranes.

Measurements from fixed tissues revealed that EGF binds rapidly to the media-exposed membranes on the surface of the cluster but is excluded from lateral membranes and from cells located within the interior of the cluster (Farahani et al., 2021). To investigate the dynamics of this spatial pattern, we monitored EGFR signaling using two live-cell biosensors: EGFR pYtag and a kinase translocation reporter for downstream signaling through Erk (ErkKTR) (Regot et al., 2014). We treated MCF10A cells cultured on soft substrata with EGF and observed a radially directed wave of EGFR and Erk signaling: activation first appeared in cells at the periphery of the clusters before appearing in cells at the interior (**Figures 2F; Video S2**). In cells at the periphery of clusters, we found that ZtSH2 was first enriched at the media-exposed membrane before localizing to lateral membranes (**Figure 2G**), highlighting differences in receptor-level signaling between membrane sub-compartments. Quantifying EGFR and Erk responses confirmed our qualitative observations, revealing a 2-4 min delay in EGFR and Erk signaling between the periphery and interior of clusters (**Figures 2H and 2I**). Notably, Erk responses also trailed those of EGFR by ∼4 min, consistent with the delay in signal transmission previously observed from Ras to Erk (**Figure 2J**) (Toettcher et al., 2013). These data support a model in which ligand-receptor interactions are limited at cell-cell contacts, producing an inward-traveling wave of RTK activation and downstream signaling. More broadly, our data demonstrate that pYtags can be used to reveal RTK signaling dynamics at seconds-scale resolution and over both subcellular and multicellular length scales.

### pYtags distinguish ligand- and dose-dependent signaling dynamics

We next applied the EGFR pYtag to study an unresolved feature of RTK signaling dynamics. While it has long been known that different RTKs regulate cellular processes by altering signaling dynamics (Freed et al., 2017; Johnson and Toettcher, 2019; Marshall, 1995; Santos et al., 2007), recent evidence suggests that different ligands alter signaling dynamics through the same receptor (Freed et al., 2017). For instance, high-affinity EGFR ligands such as EGF produce long-lived EGFR dimers that are internalized and degraded, whereas low-affinity ligands such as epiregulin (EREG) and epigen (EPGN) induce shorter-lived EGFR dimers that prolong signaling, leading to distinct outcomes in cell fate (Freed et al., 2017; Roepstorff et al., 2009). How these different ligands might alter minutes-timescale EGFR signaling dynamics in single cells remains unknown.

To answer this question, we treated EGFR pYtag-expressing NIH 3T3 cells with varying doses of either EGF or one of two low-affinity ligands, EREG or EPGN. We found that the kinetics of EGFR activation varied substantially with the dose of EGF, from a gradual rise in activity at lower doses (0.2-2 ng/mL) to a more rapid, biphasic response at higher doses (5-100 ng/mL) (**Figure 3A, left panel**). In contrast, treatment with the low-affinity ligands EREG and EPGN led to a rapid rise in EGFR activity followed by a plateau within minutes (**Figure 3A, middle and right panels**). We also observed a small but reproducible transient peak of EGFR activation shortly after treatment with EREG and EPGN but not with EGF, suggestive of weak negative feedback after stimulation with low-affinity ligands (**Figure 3A**). These results demonstrate that different ligands produce profoundly different signaling responses in the minutes immediately after EGFR stimulation. They also reveal an unusual property of the low-affinity RTK ligands EREG and EPGN: altering their concentrations has a strong effect on the amplitude, but not the kinetics, of EGFR activation.

**Figure 3.**
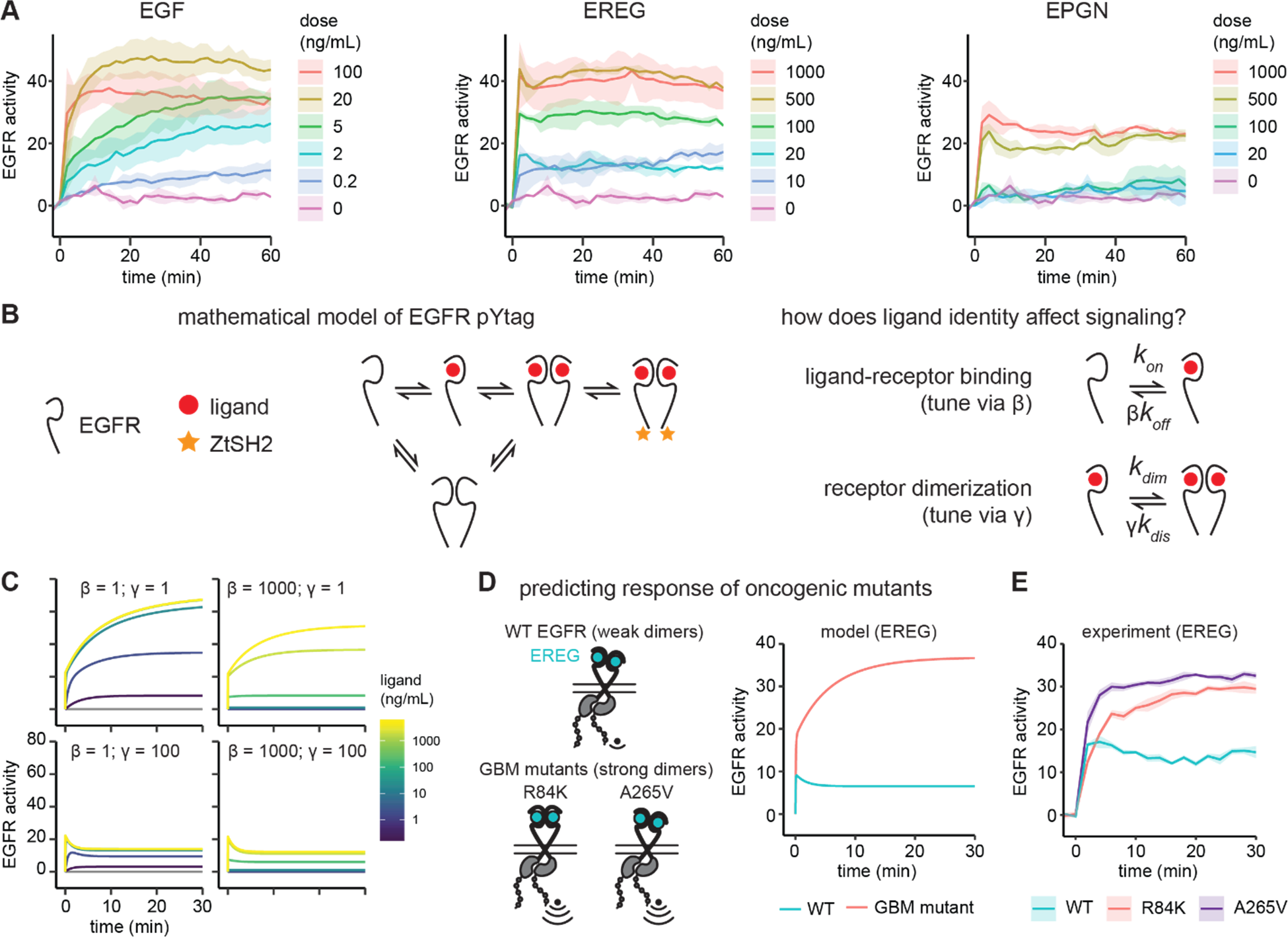
EGFR pYtag reveals dose- and ligand-dependent signaling dynamics. (A**) Mean ±** S.D. responses of EGFR pYtag-expressing NIH 3T3 cells to varying doses of EGF, epiregulin (EREG), and epigen (EPGN). The same 0 ng/mL control was used for each ligand. *n* = 2 independent experiments. (**B**) Dose-response profiles from (A) were analyzed using a mathematical model of EGFR pYtag. (**C**) Simulations of EGFR pYtag responses to ligand of varying doses, for different values of β and γ. (**D**) GBM-associated mutants of EGFR that form strong EREG-bound dimers were predicted to exhibit stronger pYtag responses to 20 ng/mL EREG compared to WT EGFR. (**E**) Mean ± S.D. pYtag response of WT and GBM-associated mutant EGFRs in NIH 3T3 cells after EREG treatment (20 ng/mL). *n* = 3 independent experiments.

We constructed a mathematical model of the EGFR pYtag based on mass-action kinetics to better understand these ligand-dependent EGFR signaling responses (**Figure 3B**). In our model, ligand-free EGFR monomers bind to ligands, dimerize, undergo autophosphorylation and, lastly, recruit ZtSH2 to ligand-bound dimers. We also modeled EGFR dimerization in the absence of ligand, based on several observations of these inactive dimers (Bessman et al., 2014; Moriki et al., 2001; Saffarian et al., 2007; Yu et al., 2002). In our model, ligand-free dimers may also bind to ligand and recruit ZtSH2, bypassing the intermediate step of transitioning from monomer to dimer. Notably, we found that ligand-free EGFR dimers were necessary to reproduce the biphasic signaling dynamics we observed after stimulation with EGF (**Figure S2**). We assumed that different EGFR ligands could alter two rate constants in the model (**Figure 3B**). First, high- and low-affinity ligands are expected to bind to their receptor with different affinities, which we modeled by varying the dissociation rate of ligand-receptor complexes using a scaling parameter β. Second, low-affinity ligands have been reported to produce structurally different EGFR dimers compared to high-affinity ligands, thereby reducing the dimerization affinity of ligand-bound receptors (Freed et al., 2017; Hu et al., 2022); we modeled this dimerization affinity by varying the dissociation rate of ligand-bound dimers using a scaling parameter γ.

To define which ligand-specific interactions correlate with differential signaling dynamics (**Figure 3A**), we systematically varied β and γ while simultaneously varying ligand doses. Setting both β and γ to 1 reproduced the dynamics of EGFR signaling in response to EGF (**Figures 3A and 3C**). We then decreased ligand binding affinity by increasing β, which increased the dose of ligand required for activation but failed to qualitatively alter the shape of the response curves as we had observed experimentally (**Figures 3A and 3C**). In contrast, decreasing the dimerization affinity of ligand-bound receptors by increasing γ was sufficient to reproduce many of the features that varied with ligand identity (**Figures 3A and 3C**). At low values of γ, we observed an EGF-like response consisting of a biphasic and gradual approach to steady-state (**Figure 3C**), due to the initial activation of ligand-free dimers, and subsequent dimerization and activation of the remaining monomeric receptors. In contrast, high values of γ produced an EREG/EPGN-like initial peak of receptor activation followed by a plateau (**Figure 3C**), due to rapid activation of ligand-free dimers but less additional dimerization of ligand-bound monomeric receptors. In sum, our model predicts that the EGFR signaling dynamics produced by high- and low-affinity ligands arise from altered dimerization affinities of ligand-bound receptors.

To validate the prediction of our model, we experimentally altered the dimerization affinity of EGFR and monitored signaling responses using pYtags. We turned to glioblastoma multiform (GBM)-associated mutations in the extracellular domain of EGFR (R84K and A265V point mutations), which were previously shown to increase the dimerization affinity of EREG- and EPGN-bound receptors (Hu et al., 2022). We first modeled these GBM-associated mutations in EGFR by decreasing β and γ by 6-fold and 650-fold, respectively (Hu et al., 2022). These modifications led to the prediction that 20 ng/mL EREG would induce a stronger, more gradual signaling response in GBM-associated mutants compared to the wild-type receptor (**Figure 3D**). We then generated NIH 3T3 cell lines expressing pYtag biosensors of EGFR variants harboring either the R84K or A265V mutation. Treating these cells with 20 ng/mL EREG induced signaling responses that closely matched the predictions of our model: cells expressing WT EGFR exhibited a transient peak of activation and rapid plateau, while cells expressing GBM-associated mutants exhibited a stronger response that gradually increased over time (**Figures 3D and 3E**). Our data reveal ligand-dependent EGFR signaling dynamics at unprecedented temporal resolution and point to the dimerization affinity of ligand-bound receptors as a key parameter that governs these responses. More broadly, combining pYtags with mathematical modeling can shed mechanistic insight into the signaling dynamics of RTKs.

### pYtags report the activity of distinct RTKs in heterodimeric complexes

One advantage of the pYtag approach is its modularity: ITAMs can in principle be fused to the C-termini of many different RTKs and recruit ZtSH2 upon stimulation. We thus tested whether pYtags could be adapted to monitor the activity of receptors other than EGFR. We designed pYtag biosensors for two additional RTKs: 1) ErbB2, a ligandless member of the ErbB family that signals by heterodimerizing with ligand-bound EGFR (Lemmon and Schlessinger, 2010); 2) fibroblast growth factor receptor 1 (FGFR1), a non-ErbB family RTK. ErbB2 is a particularly challenging RTK to study because it can only be activated in conjunction with an additional RTK (e.g., EGFR), so Grb2-based biosensors (**Figure S1**) would be unable to distinguish between the activity of EGFR and ErbB2. Starting from NIH 3T3 cells expressing iRFP-ZtSH2, we generated cell lines expressing either an ErbB2 pYtag (ErbB2-CD3ε- FusionRed), an FGFR1 pYtag (FGFR1-CD3ε-FusionRed), or ITAM-less variants of each. For the ErbB2 case, we further transduced cells with EGFR-Citrine to express both required components of a functional receptor heterodimer (**Figure 4A**).

**Figure 4.**
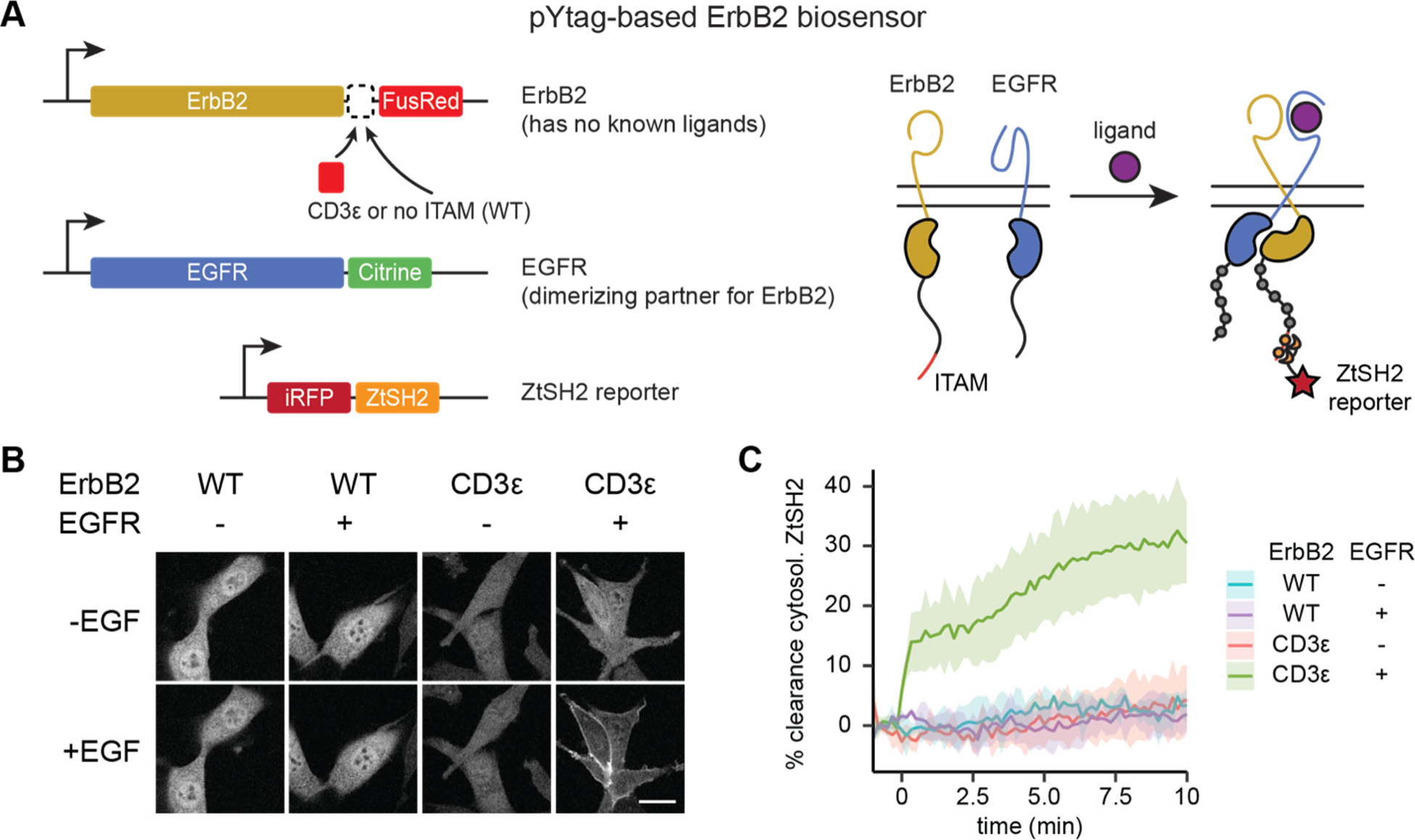
Monitoring distinct RTKs in heterodimeric complexes. (**A**) In order to signal, the ligandless ErbB2 must heterodimerize with a ligand-binding member of the ErbB family. ITAM appendage to ErbB2 enables measurements of the receptor’s activity despite the co-activation of EGFR. (**B**) Representative images of NIH 3T3 cells treated with EGF (100 ng/mL). Scale bar, 20 µm. (**C**) Mean ± S.D. clearance of cytosolic ZtSH2 after treatment with EGF (100 ng/mL). *n* = 3 independent experiments.

We found that pYtags generalized well to both ErbB2 and FGFR1. As with the EGFR pYtag, we quantified the responses of the ErbB2 and FGFR1 pYtags by measuring the clearance of ZtSH2 from the cytosol in response to treatment with activating ligands. Stimulating ErbB2 pYtag-expressing cells with EGF led to clearance of ZtSH2 from the cytosol, a response that required both the presence of ITAMs on ErbB2 and co-expression of EGFR (**Figures 4B and 4C**). Similarly, stimulating FGFR1 pYtag-expressing cells with FGF4 induced cytosolic clearance of ZtSH2, which did not take place in cells that expressed an ITAM-less FGFR1 (**Figure S3**). Both FGFR1-FusionRed and FGFR1-CD3ε-FusionRed were also prominently clustered in internal compartments that recruited ZtSH2 in an ITAM-dependent manner, suggesting a high degree of basal signaling from these internal compartments that could be due to ectopic expression of the receptor (**Figure S3A**). The pYtag approach is indeed modular, and this strategy can be readily adapted for monitoring the activation of distinct RTKs in individual cells.

### Multiplexing RTK biosensors using orthogonal pYtags

Our pYtag design strategy relies on the high selectivity that can be produced by multivalent association between pairs of phosphotyrosine motifs and tSH2 domains. We thus hypothesized that two orthogonal pYtags could be deployed to monitor the activation of distinct RTKs in the same cell. To this end, we took advantage of another phosphotyrosine/tSH2 interaction pair from immune-specific signaling proteins to build a pYtag that functions orthogonally to the above-described CD3ε/ZtSH2 system. Following its binding to phosphorylated ITAMs on the TCR, ZAP70 phosphorylates non-ITAM tyrosine residues on the scaffold protein SLP76, which subsequently recruit multiple signaling components via SH2- mediated interactions (Chakraborty and Weiss, 2014; Weiss and Littman, 1994). Because immune signaling requires these events to be discreet, sequential, and to operate in proximity to each other, ZtSH2- and SLP76-recruited SH2s must have orthogonal binding specificities. We exploited this feature to engineer a second pYtag. We identified two tyrosine residues of SLP76, Y128 and Y145, which when phosphorylated bind to SH2 domains from the guanine nucleotide exchange factor Vav and the kinase ITK, respectively (**Figure 5A**) (Koretzky et al., 2006; Raab et al., 1997; Su et al., 1999). To create a synthetic tSH2 that could bind tightly to the Y128/Y145 motif, we fused the SH2 domains of Vav and ITK together with a 10 bp glycine-serine linker to create VISH2. We then characterized the suitability of the SLP76/VISH2 system for monitoring RTK activity and quantified its crosstalk with the CD3ε/ZtSH2 system.

**Figure 5.**
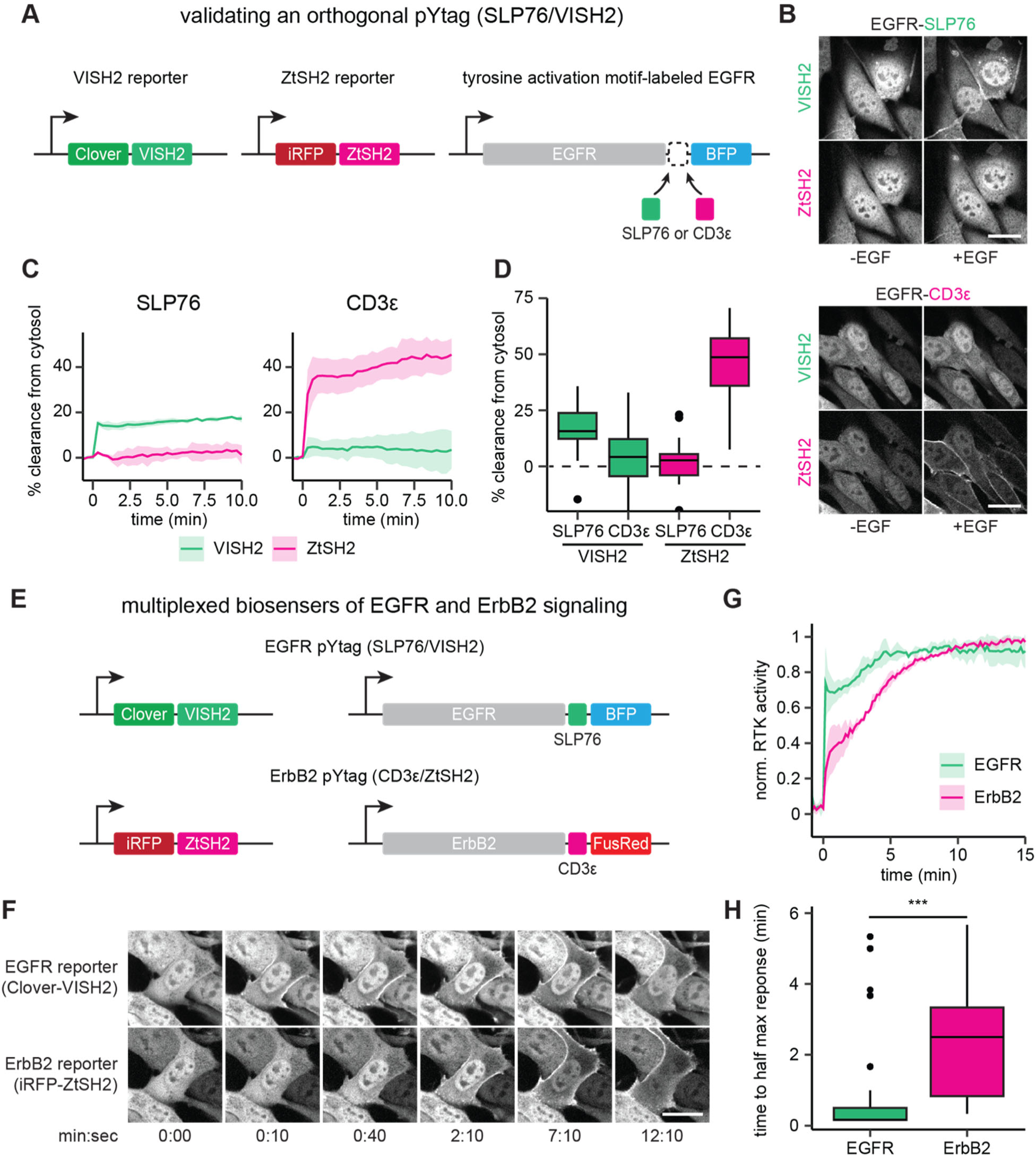
Orthogonal pYtags enable multiplexed RTK biosensing. (**A**) To assess the performance of the VISH2/SLP76 system as a pYtag-based biosensor, VISH2 and ZtSH2 reporters were co-expressed in NIH 3T3 cells along with either SLP76- or CD3ε-labeled EGFR. (**B**) NIH 3T3 cells co-expressing VISH2 and ZtSH2 reporters before and 3 min after treatment with EGF (100 ng/mL). Scale bars, 20 µm. (**C**) Mean ± S.D. clearances of VISH2 and ZtSH2 from the cytosol, expressed with either SLP76- or CD3ε-labeled EGFR and stimulated with EGF (100 ng/mL). *n* = 3 independent experiments. (**D**) Response of VISH2 and ZtSH2 reporters 10 min after EGF treatment in (C). *n* > 30 cells from 3 independent experiments. (**E**) Orthogonal pYtags can be multiplexed to monitor the activity of multiple RTKs in the same cell. (**F**) Images of cells expressing orthogonal reporters for EGFR and ErbB2, treated with EGF (100 ng/mL). Scale bar, 20 µm. (**G**) Mean ± S.D. trajectories for EGFR and ErbB2 activity using multiplexed pYtags. For each reporter, the mean response was normalized to its minimum and maximum measured values. *n* = 3 independent experiments. (**H**) Time to half maximal response for individual cells from (G). Lines denote mean values, boxes denote 25-75^th^ percentiles, and whiskers denote minima and maxima. *n* > 30 cells from 3 independent experiments. *** *p* < 0.001 by Kolmogorov-Smirnov test. See also Videos S3 and S4.

To determine whether the SLP76/VISH2 system functions as a pYtag biosensor in an orthogonal manner to CD3ε/ZtSH2, we generated NIH 3T3 cells that stably co-expressed Clover-VISH2 and iRFP-ZtSH2; we then further transduced these cells with a variant of EGFR labeled with either three repeats of the E123-E153 region of SLP76 (EGFR-SLP76-BFP) or three repeats of the CD3ε ITAM (EGFR-CD3ε-BFP) (**Figure 5A**). Stimulating the two cell lines with EGF and monitoring the localization of VISH2 and ZtSH2 revealed that cells expressing SLP76- labeled EGFR exhibited clearance of VISH2 from the cytosol but no crosstalk with ZtSH2 (**Figures 5B-5D; Video S3**). Conversely, cells expressing CD3ε-labeled EGFR exhibited strong clearance of ZtSH2 from the cytosol with minimal crosstalk with VISH2 (**Figures 5B-5D; Video S3**). The VISH2/SLP76 and CD3ε/ZtSH2 pYtags thus operate independently of one another to report the activity of EGFR.

Having characterized an orthogonal pair of pYtags, we hypothesized that these biosensors could be used to simultaneously monitor the activation of both EGFR and ErbB2. To test this hypothesis, we generated NIH 3T3 cells that co-expressed pYtags for EGFR (Clover-VISH2; EGFR-SLP76-BFP) and ErbB2 (iRFP-ZtSH2; ErbB2-CD3ε-FusionRed) (**Figure 5E**). We treated cells with 100 ng/mL EGF and observed that both VISH2 and ZtSH2 reporters cleared from the cytosol, as would be expected from the activation of EGFR and ErbB2 heterodimers. However, the biosensors revealed markedly different dynamics of receptor activation (**Figures 5F-5H; Video S4**). VISH2 cleared from the cytosol almost immediately (<30 sec), indicating rapid phosphorylation of the EGFR C-terminal tail. In contrast, ZtSH2 exhibited a more gradual clearance from the cytosol (∼3 min), suggesting a delay in ErbB2 phosphorylation. To verify that these dynamics are not due to the identity of the pYtag used to monitor each receptor or an artifact of the cell line expressing multiplexed biosensors, we compared the dynamics of EGFR and ErbB2 activation across all experiments that used 100 ng/mL EGF, regardless of the cell line or pYtag used (**Figure S4**). Although the absolute magnitude of tSH2 biosensor clearance varied between cell lines (likely due to differences in affinity and expression level), the dynamics of EGFR and ErbB2 activation were highly reproducible in all cases, irrespective of the cell line or pYtag (**Figures S4B-S4C**). These data demonstrate that pYtag measurements faithfully reflect the activation of the RTK on which they report. Our results suggest that EGFR and ErbB2 are activated in distinct phases: EGFR is activated first, within seconds of stimulation, and then both EGFR and ErbB2 are further activated over the subsequent minutes.

### pYtags can monitor the activity of endogenous RTKs

One drawback to the pYtag approach is its reliance on ectopically expressing an ITAM- labeled receptor of interest. Increasing the expression levels of RTKs can alter signaling dynamics, downstream pathway engagement, and cell fate outcomes (Dikic et al., 1994; Traverse et al., 1994). Conversely, the low expression levels of endogenous RTKs may only drive weak recruitment of tSH2, making biosensing by live-cell microscopy difficult. We thus asked whether pYtags could be adapted to monitor the activity of endogenously expressed RTKs.

We used CRISPR/Cas9-based genome editing to label the C-terminus of endogenous EGFR with three repeats of the CD3ε ITAM and a fluorescent protein via homology-directed repair (HDR) in HEK 293T cells (**Figure 6A**). We chose mNeonGreen as our fluorescent protein label because of its exceptional brightness, which aids detection at low levels of endogenous expression (Shaner et al., 2013). We sorted an mNeonGreen-expressing clonal cell line and confirmed proper labeling of EGFR with CD3ε-mNeonGreen by PCR of genomic DNA (**Figure 6B**) and immunoblotting (**Figure 6C**). We then transduced both parental 293T and EGFR-CD3ε- mNeonGreen-expressing (knock-in 293T) cells with an mScarlet-labeled ZtSH2 to take advantage of this fluorescent protein’s high brightness for detecting subcellular ZtSH2 localization even at low expression levels (Bindels et al., 2017). We then monitored ZtSH2 localization after treatment with EGF in both parental and knock-in cells. Knock-in 293T cells exhibited rapid clearance of ZtSH2 from the cytosol following EGF treatment, whereas no ZtSH2 response was observed in parental cells (**Figures 6D-6F**). We observed striking differences in the dynamics of endogenous receptor activation compared to experiments in which we ectopically expressed EGFR: endogenous EGFR exhibited a decrease in activity within minutes after EGF stimulation (**Figure 6D; Video S5**) whereas membrane recruitment was sustained for at least 60 min for ectopically expressed EGFR (**Figure 3A**). We also observed rapid and near-complete internalization of endogenous EGFR from the cell membrane, with some internalized vesicles retaining residual ZtSH2 labeling (**Figure 6D, right-most panels**).

**Figure 6.**
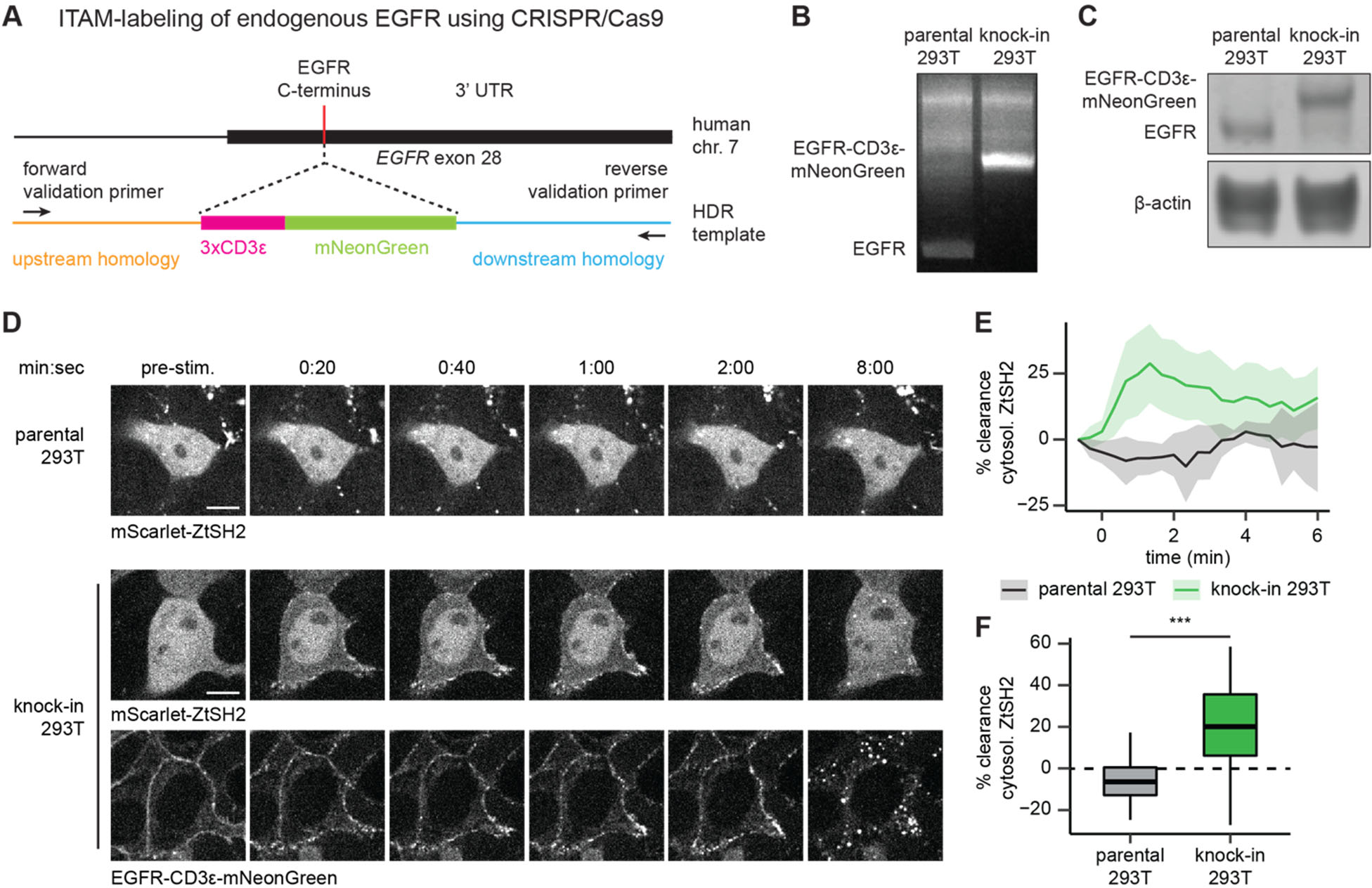
pYtags can monitor the activity of endogenous RTKs. (**A**) Schematic of *EGFR* locus containing the C-terminus of EGFR, where CRISPR/Cas9 was used to label the receptor with CD3ε-mNeonGreen via homology-directed repair. (**B**) PCR of genomic DNA from parental or knock-in 293T cells. Validation primers targeting homology regions upstream and downstream of the CD3ε-mNeonGreen insert are labeled by black arrows in (A). (**C**) Immunoblots of EGFR in parental or knock-in 293T cells. (**D**) Images of parental or knock-in 293T cells treated with EGF (100 ng/mL). mScarlet-ZtSH2 images show averages of two successive frames to decrease background noise; full raw movie is included as Video S5. Scale bar, 10 µm. (**E**) Mean ± S.D. clearance of ZtSH2 from the cytosol following treatment with EGF (100 ng/mL). parental 293T, *n* = 3 independent experiments; knock-in 293T, *n* = 4 independent experiments. (**F**) Clearance of ZtSH2 from the cytosol 1 min after treatment with EGF in (E). Lines denote mean values, boxes denote 25-75^th^ percentiles, and whiskers denote minima and maxima. parental 293T, *n* = 23 cells from 3 independent experiments; knock-in 293T, *n* = 46 cells from 4 independent experiments. *** *p* < 0.001 by Kolmogorov-Smirnov test. See also Video S5.

We therefore conclude that pYtags can be used to monitor the activation state of endogenously expressed RTKs, opening the door to single-cell studies of RTK signaling dynamics in minimally perturbed contexts.

## Discussion

Here we describe pYtags, a biosensing strategy for monitoring the activity of a specific RTK in living cells. pYtags rely on tyrosine activation motifs that are selectively bound by corresponding tSH2 domains to minimize interactions with endogenous phosphotyrosine motifs and SH2 domains. This design principle confers selectivity to certain steps in T cell signaling, such as the recruitment of the kinase ZAP70 to phosphorylated ITAMs within activated TCRs. We exploit this property of immune signaling proteins to build two orthogonal pYtags: the first based on ITAMs from the TCR and the tSH2 domain of ZAP70, and the second based on the tyrosine-containing scaffold protein SLP76 and its SH2-containing binding partners Vav and ITK. We show that pYtags can be applied to multiple RTKs (EGFR, ErbB2, and FGFR1), providing a robust strategy to monitor receptor-level signaling in living cells.

pYtags quantitatively report on RTK activity with seconds-scale precision and can be applied to monitor signaling at both subcellular and multicellular length scales (**Figure 2**).

Responses are highly reproducible across multiple experiments and cell lines, even when the same receptor is monitored using different pYtag variants (**Figure S4**). This quantitative precision enabled us to identify differences in EGFR signaling dynamics induced by high- or low-affinity ligands (**Figure 3**), and to observe distinct signaling dynamics for two ErbB-family receptors (**Figure 5**). We used pYtag measurements to inform a mathematical model that, along with subsequent experimental validation, suggests that the dimerization affinity of ligand-bound receptors controls RTK signaling dynamics (**Figure 3**). Our data suggest that pYtags represent a broadly applicable biosensing strategy to monitor the activity of any RTK of interest and can reveal new mechanistic insights for even the most well-studied RTKs.

pYtags have several advantages over previously reported biosensors of RTK activity. First, pYtags are modular: they can be applied to multiple RTKs without modification, unlike approaches that seek to identify phosphotyrosine/SH2 interactions present on endogenous RTKs (Tiruthani et al., 2019). Second, pYtags are specific, reporting only on the activity of the RTK labeled by the tyrosine activation motif. Third, pYtags can be multiplexed: pYtags require one fewer fluorescent protein compared to FRET-based biosensors, and orthogonal variants of pYtags can be used to monitor distinct RTKs in the same cell. The human genome encodes at least 58 RTKs (Lemmon and Schlessinger, 2010), most of which remain poorly characterized with respect to their spatial organization and signaling dynamics. Multiplexing pYtags could open the door to systematically characterizing how different combinations of RTKs and cognate ligands influence signal processing, as has been performed for the bone morphogenetic protein receptor family (Antebi et al., 2017; Klumpe et al., 2022; Su et al., 2022).

Despite the advantages of pYtags, there are several challenges that might limit the adoption of this strategy. First, pYtags require a tyrosine activation motif to be appended to an RTK of interest, necessitating either ectopic expression of the labeled receptor or direct genome editing of the endogenous receptor. Although the increasingly powerful toolbox available for CRISPR-based genome editing lessens the burden associated with this strategy, endogenous tagging remains technically challenging and must be considered when engineering biosensor-expressing cell lines or organisms. A second limitation of pYtags is its reliance on phosphotyrosine/tSH2 motifs from immune signaling proteins. As a result, pYtags are currently restricted to non-immune cells that do not express the corresponding endogenous proteins. It would be useful to further extend the approach to *de novo* designed substrate/binder pairs to broaden their applicability to additional cellular contexts.

We anticipate many future applications of the pYtag biosensing strategy. Live-cell biosensors of Erk signaling have revealed complex spatiotemporal signaling patterns – pulses, oscillations, and traveling waves – that appear to depend on the activation of certain RTKs (De Simone et al., 2021; Hino et al., 2020; Hiratsuka et al., 2015; Pokrass et al., 2020; Regot et al., 2014; Simon et al., 2020). Yet it remains unclear whether pulses of Erk activity require pulsatile activation of upstream RTKs. pYtags also open the door to quantitatively characterizing long-range ligand gradients that are thought to underlie processes such as collective cell migration or morphogen signaling during development (Hino et al., 2020; Sprenger and Nüsslein-Volhard, 1992). Finally, we expect that the design principle of pYtags could be applied beyond RTK biosensors, such as for monitoring the signaling of non-receptor tyrosine kinases or engineering synthetic receptors with user-defined response programs.

## Acknowledgments

We thank all members of the Bashor, Nelson, and Toettcher labs for their insights and comments, and Gary Laevsky and Sha Wang from the Princeton Nikon Imaging Facility for assistance with microscopy. This work was supported by NSF CAREER Award 1750663 and a Vallee Scholar award (to J.E.T.); NIH grants R01HL164861 and R01HD099030 (to C.M.N.); a Schmidt Transformative Technology Award (to C.M.N. and J.E.T.); NIH training grant T32GM007388 (to E.V.M.); NIH grant R01EB032272 and ONR grant N00014-21-1-4006 (to C.J.B.). P.E.F. was supported in part by the NSF Graduate Research Fellowship Program.

## Author contributions

Conceptualization, P.E.F, X.Y, C.J.B, C.M.N, and J.E.T; experimental investigation, P.E.F, X.Y., E.V.M, and K.A.F; mathematical modeling, P.E.F; data analysis, P.E.F and E.V.M.; writing, P.E.F, C.M.N., and J.E.T, with input from all authors; funding acquisition and supervision, C.J.B, C.M.N, and J.E.T.

## Resource details

### Lead contact

Further information and requests for resources and reagents should be directed and will be fulfilled by the lead contact, Jared Toettcher.

### Materials availability

There are no restrictions on material availability. Plasmids are available from Addgene (www.addgene.org/Jared_Toettcher); all cell lines produced and plasmids unavailable on Addgene will be made available upon request.

### Data and code availability

- There are no restrictions on data availability. Source data generated or analyzed during this study, as well as Python and R scripts for data analysis and mathematical modeling, are available on the laboratory GitHub page (https://github.com/toettchlab).

## Key resources table

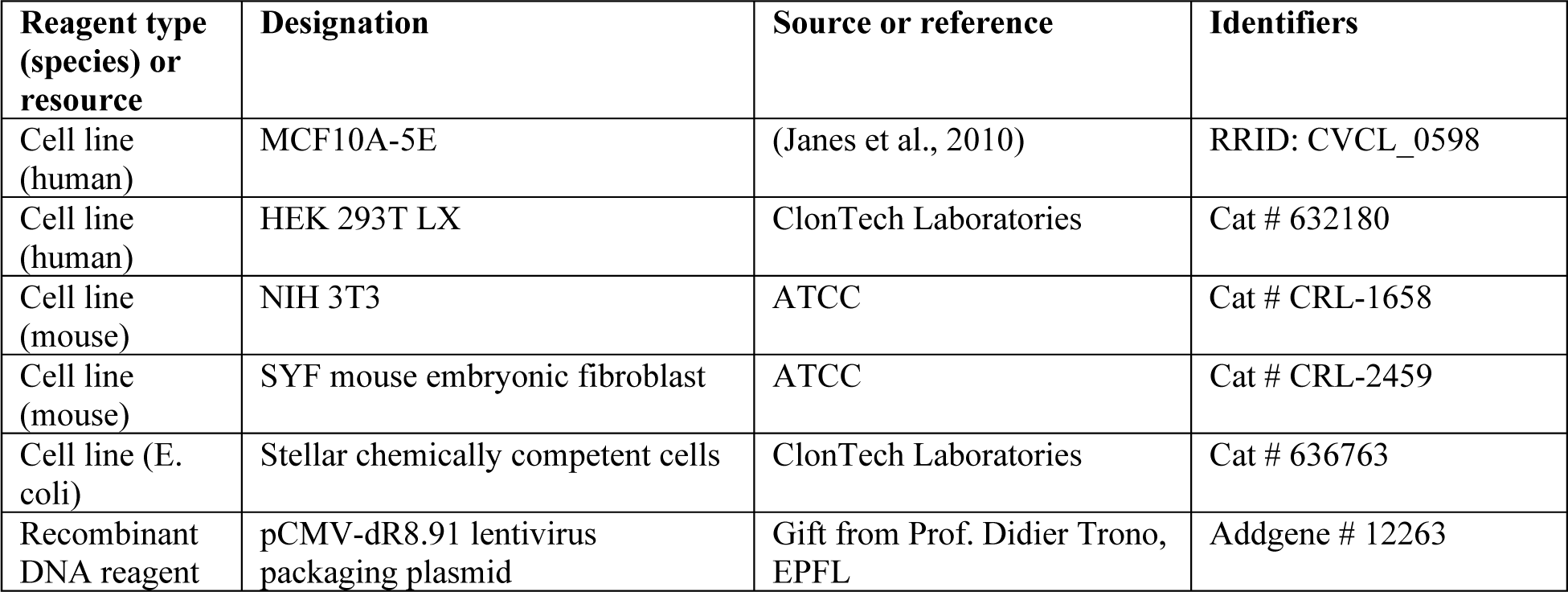
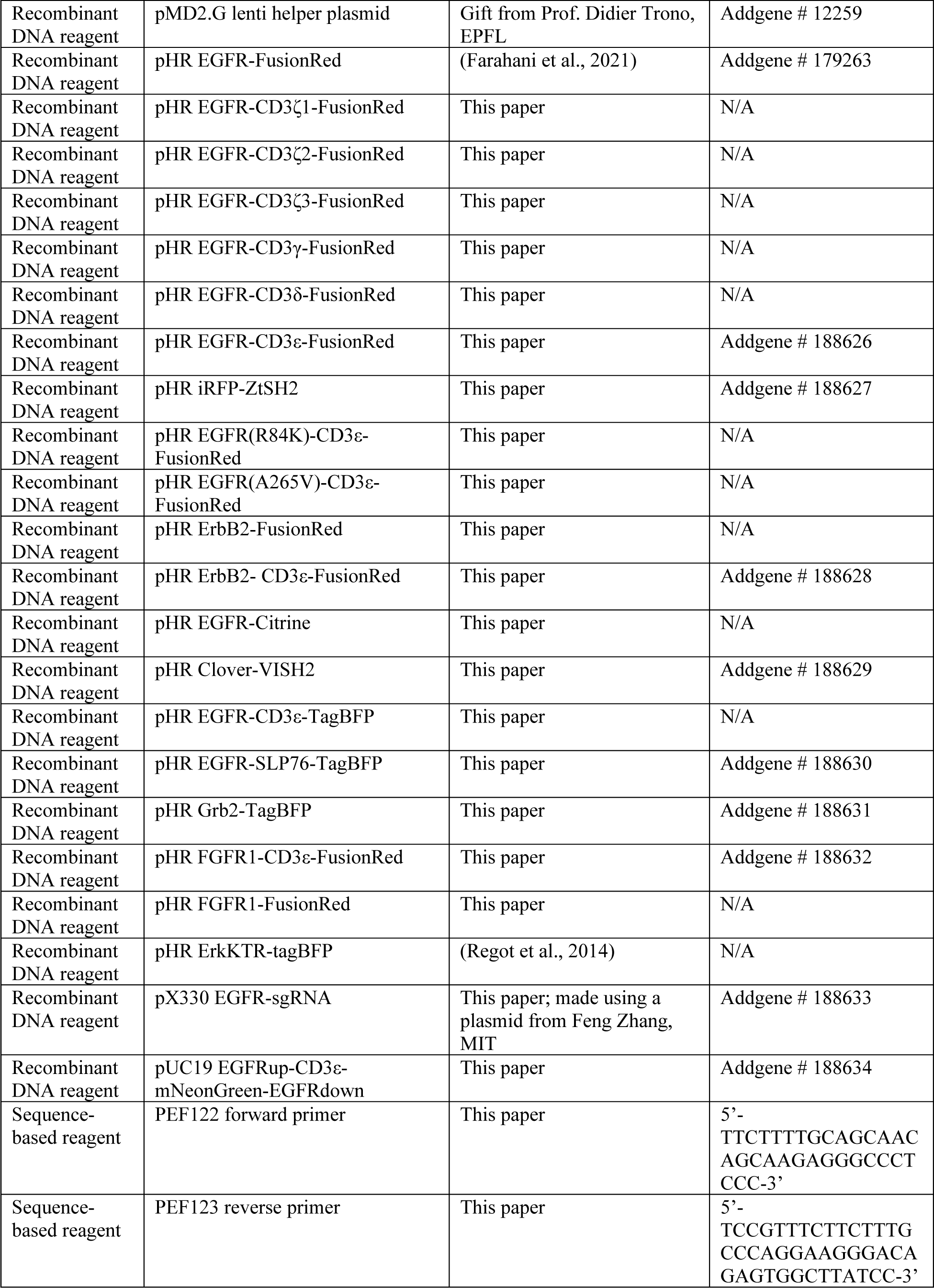
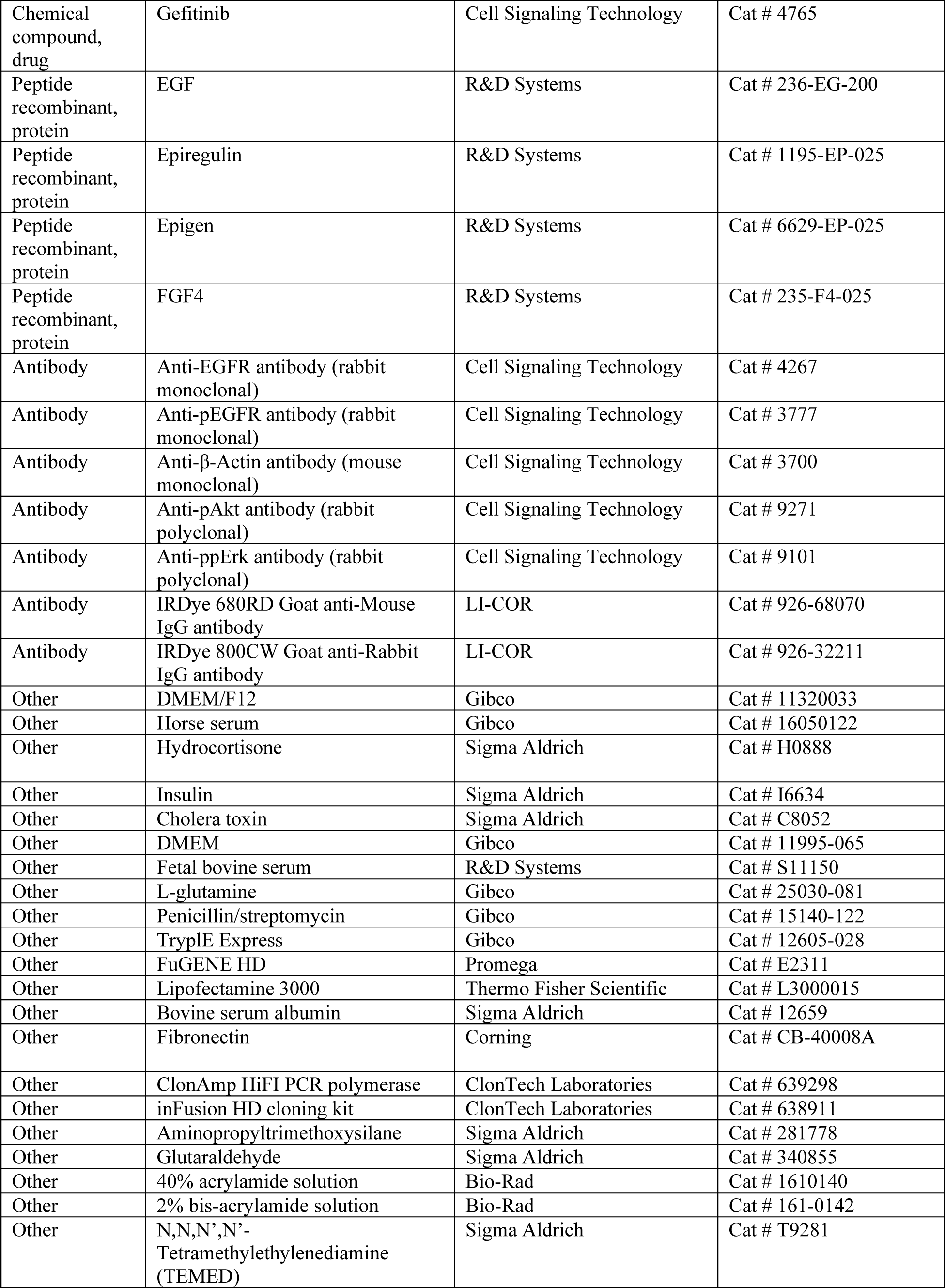
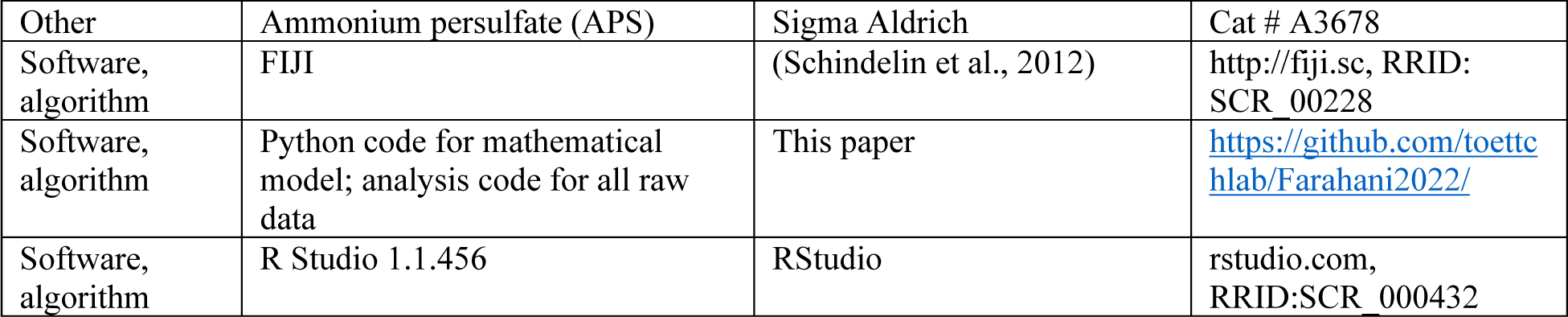

## Methods

### Plasmid construction

All constructs were cloned into the pHR lentiviral expression plasmid using inFusion cloning. Linear DNA fragments were produced by PCR using HiFi polymerase (Takara Bio), followed by treatment with DpnI to remove template DNA. PCR products were then isolated through gel electrophoresis and purified using the Nucleospin gel purification kit (Takara Bio). Linear DNA fragments were then ligated using inFusion assembly and amplified in Stellar competent *E. coli* (Takara Bio). Plasmids were purified by miniprep (Qiagen) and verified by either Sanger sequencing (Genewiz) or whole-plasmid sequencing (Plasmidsaurus).

### Cell line generation

Constructs were stably expressed in cells using lentiviral transduction. First, lentivirus was produced by co-transfecting HEK 293T LX cells with pCMV-dR8.91, pMD2.G, and the expression plasmid of interest. 48 h later, viral supernatants were collected and passed through a 0.45 µm filter. Cells were seeded at ∼30% confluency and transduced with lentivirus 24 h later.

24 h post-seeding, culture medium was replaced with medium containing 10 µg/mL polybrene and 200-300 µL viral supernatant was added to cells. Cells were then cultured in virus-containing medium for 48 h. Populations of cells co-expressing each construct were isolated using fluorescence-activated cell sorting on a Sony SH800S cell sorter. Bulk-sorted populations were collected for all experiments, except for those using MCF10A cells and CRISPR/Cas9- edited HEK 293T cells, for which clonal lines were generated.

### CRISPR/Cas9-based gene editing

HEK 293T cells with CD3ε-mNeonGreen inserted at the endogenous *EGFR* locus were generated by transfecting cells with 1) a pX330 plasmid containing a human codon-optimized SpCas9 and guide RNA targeting *EGFR* (pX330 EGFR-sgRNA) and 2) a homology-directed repair template comprised of three repeats of the CD3ε ITAM and mNeonGreen flanked by 800 bp of *EGFR* homology regions (pUC19 EGFRup-CD3ε-mNeonGreen-EGFRdown) (Ran et al., 2013). HEK 293T cells were first seeded at 130,000 cells/well in a 24-well plate. 24 h later, cells were transfected with 330 ng pX330 EGFR-sgRNA and 170 ng pUC19 EGFRup-CD3ε- mNeonGreen-EGFRdown using Lipofectamine 3000 (Thermo Fisher Scientific) and left to culture for another 48 h. Clonal cell lines were then isolated using fluorescence-activated cell sorting on a Sony SH800S cell sorter, and localization of mNeonGreen was assessed by confocal microscopy. A candidate clonal cell line exhibiting membrane-localized mNeonGreen was then used for further validation.

To verify genomic integration of CD3ε-mNeonGreen, genomic DNA was isolated from parental and knock-in 293T cells using a PureLink Genomic DNA Mini Kit (Invitrogen). PCR using HiFi polymerase was then used to amplify a region of genomic DNA using primers specific to homology regions flanking the CD3ε-mNeonGreen insertion (PEF122 and PEF123 in **Key Resources Table**). PCR products were then run through an agarose gel by electrophoresis and imaged using an Axygen Gel Documentation System. In the gel, endogenous EGFR was expected to appear as a ∼1.2 kb product, while EGFR-CD3ε-mNeonGreen was expected to appear upshifted as a ∼2.1 kb product. The expression of EGFR-CD3ε-mNeonGreen protein was verified by immunoblotting as described below.

### Cell culture

NIH 3T3 cells, HEK 293T cells, and SYF cells were cultured in DMEM (Gibco) supplemented with 10% fetal bovine serum (R&D Systems), 1% L-glutamine (Gibco), and 1% penicillin/streptomycin (Gibco). MCF10A-5E cells (Janes et al., 2010) were cultured in DMEM F12 (Gibco) supplemented with 5% horse serum (ATCC), 20 ng/mL EGF (R&D Systems), 0.5 µg/mL hydrocortisone (Corning), 100 ng/mL cholera toxin (Sigma-Aldrich), 10 µg/mL insulin (Sigma-Aldrich), and 1% penicillin/streptomycin (Gibco). All cells were maintained at 37°C and 5% CO_2_.

### Polyacrylamide substrata

To prepare polyacrylamide substrata, 1.5-mm-thick glass coverslips were pre-treated with glutaraldehyde. First, coverslips were treated with 0.1 N NaOH for 30 min, followed by rinsing with deionized water and air drying. Coverslips were then treated with 2% aminopropyltrimethoxysilane (Sigma Aldrich) in acetone for 30 min, washed three times with acetone, and left to air dry. Coverslips were treated with 0.5% glutaraldehyde (Sigma Aldrich) in PBS for 30 min, washed with deionized water, and left to air dry. Custom glass-bottom dishes were prepared by replacing the bottoms of 35 mm TCPS dishes with glutaraldehyde-treated coverslips, which were sealed using PDMS (Sigma Aldrich).

Soft (*E* ∼ 0.1 kPa) polyacrylamide substrata were made by first preparing a solution of 5% v/v acrylamide, 0.01% v/v bis-acrylamide, 0.05% v/v TEMED, and 0.05% v/v APS in deionized water. The acrylamide solution was pipetted onto a glass-bottom, glutaraldehyde-treated dish, sandwiched with an untreated coverslip, and allowed to gel for 1 h at room temperature. The untreated coverslip was then removed, leaving a polyacrylamide hydrogel attached to the glutaraldehyde-treated coverslip. To coat polyacrylamide substrata with fibronectin, substrata were first washed with ethanol, washed three times with PBS, then washed once with HEPES buffer (50 mM, pH 8.5). 1 mg/mL sulfo-SANPAH (Thermo Fisher Scientific) in deionized water was pipetted onto the hydrogel, which was then subjected to UV crosslinking (2.8 J of 365 nm light exposure over 10 min). Substrata were then rinsed once with HEPES and treated again with sulfo-SANPAH and UV crosslinking. Substrata were rinsed three times with HEPES, coated with 100 µg/mL fibronectin (Corning) in PBS, and left at 4°C overnight before seeding cells the next day.

### Preparation of samples for live-cell imaging

Cells were imaged on glass-bottom, black-walled 96-well plates (Cellvis) coated with fibronectin. Wells of 96-well plates were first incubated with 10 µg/mL fibronectin dissolved in PBS at 37°C for 30 min. Cells were seeded on glass-bottom 96-well plates at ∼20,000 cells/well 1 day prior to imaging. 4 h prior to imaging, the growth medium of cells was replaced with growth factor-free medium consisting of DMEM (Gibco) supplemented with 4.76 mg/mL HEPES and 3 mg/mL bovine serum albumin. MCF10A cells were seeded on polyacrylamide substrata at ∼400,000 cells/well 2 days prior to imaging. 24 h prior to imaging, the growth medium of cells was replaced with growth factor-free medium consisting of DMEM/F12 (Gibco) supplemented with 0.5 µg/mL hydrocortisone, 100 ng/mL cholera toxin, 3 mg/mL bovine serum albumin, and 50 µg/mL penicillin/streptomycin. To prevent evaporation of media while imaging, 50 µL of mineral oil (VWR) was pipetted onto wells prior to mounting samples on the microscope.

### Live-cell imaging

Timelapse imaging of NIH 3T3 cells, SYF cells, and HEK 293T cells was performed on a Nikon Eclipse Ti microscope with a Yokogawa CSU-X1 spinning disk, an Agilent laser module containing 405, 488, 561, and 650 nm lasers, and an iXon DU897 EMCCD camera, using 40× or 60× oil objectives. Timelapse imaging of MCF10A cells on polyacrylamide substrata was performed on a Nikon Ti2-E microscope with a CSU-W1 SoRa spinning disk, a Hamamatsu FusionBT sCMOS camera, using a 20× air objective with 2.8× magnification optics.

### Immunoblotting

Cells were lysed in ice-cold RIPA buffer (1% Triton X-100, 50 mM HEPES, 150 mM NaCl, 1.5 mM MgCl_2_, 1 mM EGTA, 100 mM NaF, 10 mM sodium pyrophosphate, 1 mM Na_3_VO_4_, 10% glycerol) supplemented with freshly prepared protease and phosphatase inhibitors. Protein levels were quantified using a Pierce BCA kit (Thermo Fisher Scientific), before being mixed with 6x Laemmli buffer/2-mercaptoethanol, heated for 5 min at 95°C, and loaded onto a 4-12% Bis-Tris gel (Invitrogen) for electrophoresis. Gels were transferred to a nitrocellulose membrane using the iBlot dry transfer system (Thermo Fisher Scientific), blocked in TBST with 5% milk for 30 min at room temperature, and incubated in primary antibody overnight at 4°C. Before imaging, membranes were washed in TBST and treated with either IRDye 680CW or 800CW secondary antibodies (LI-COR) for 1 h. Imaging was performed with the LI-COR Odyssey Infrared Imaging System. Immunoblot images were analyzed using FIJI. The signal of the target protein was normalized by the signal of β-actin, which was used as a loading control. To normalize EGFR phosphorylation levels (pEGFR) to those of total EGFR, β-actin-normalized pEGFR signals were divided by their respective β-actin-normalized EGFR signals from the same experiment.

## Mathematical modeling

### System of ordinary differential equations (ODEs)

We used a system of ODEs to model the response of EGFR pYtag to soluble ligands, in cells initially residing in medium absent of EGFR-stimulating ligands. The model captures the following events using mass-action kinetics: 1) binding between ligands and receptors; 2) dimerization and autophosphorylation of receptors, which we treat as a single event; 3) recruitment of ZtSH2 to ligand-bound EGFR dimers. Ligand binding, receptor dimerization/phosphorylation, and ZtSH2 recruitment were all treated as states that existed for individual receptors. We also assumed that soluble ligands in the culture medium above cells were in vast excess to receptors, such that the concentration of soluble ligands is held constant. Since our simulations were run on relatively short time scales (∼30 min post-stimulation), we did not consider trafficking and degradation of receptors. Recognizing that receptors could exist in every possible combination of these three states led to the following species and associated ODEs:

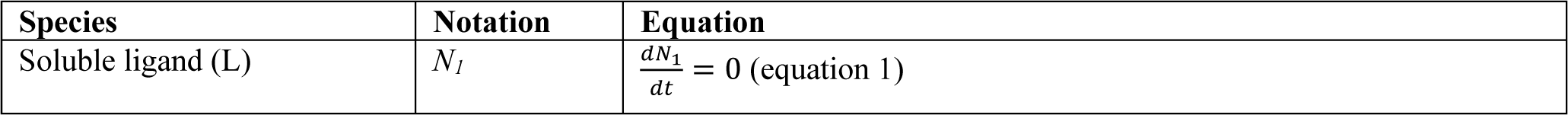
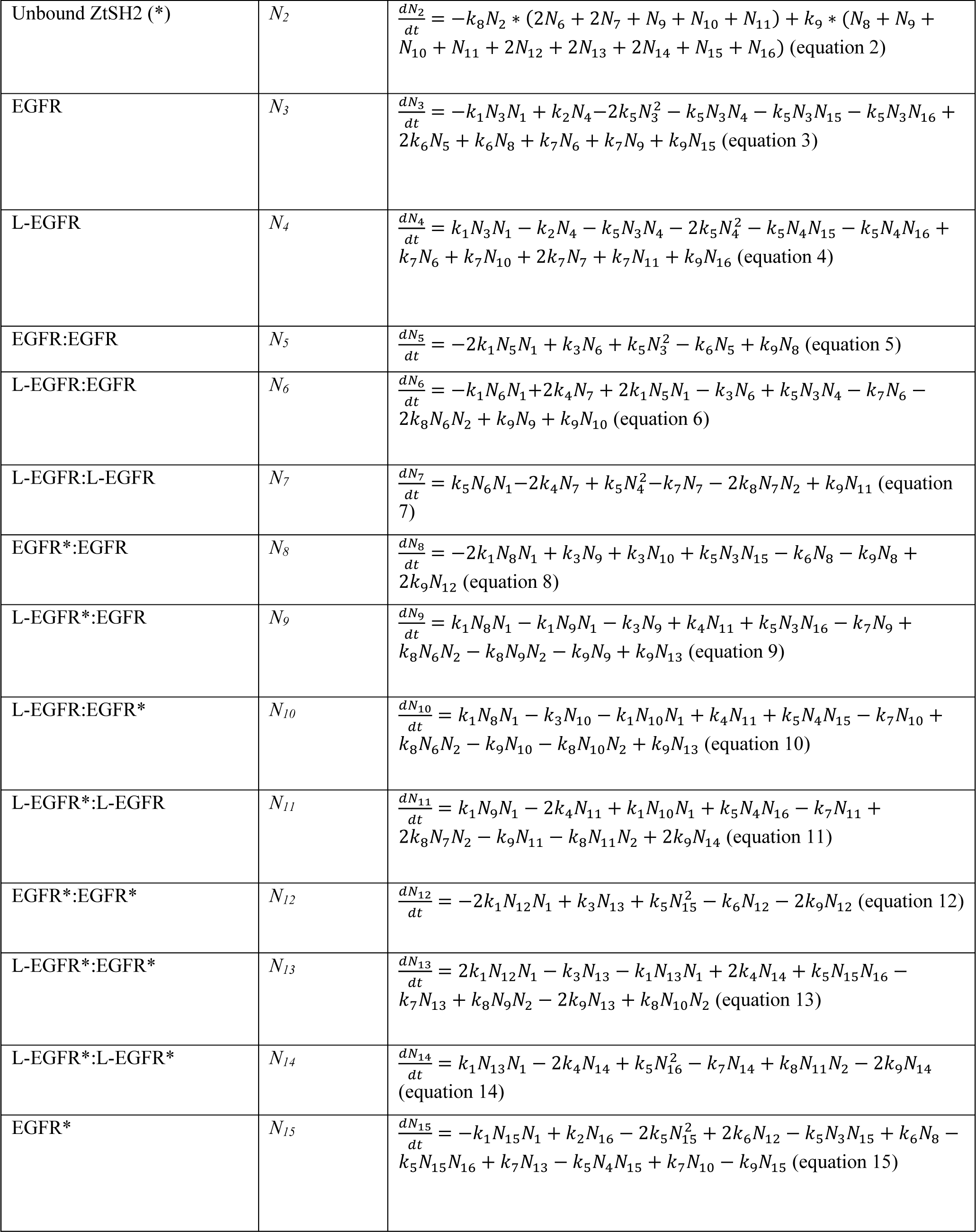
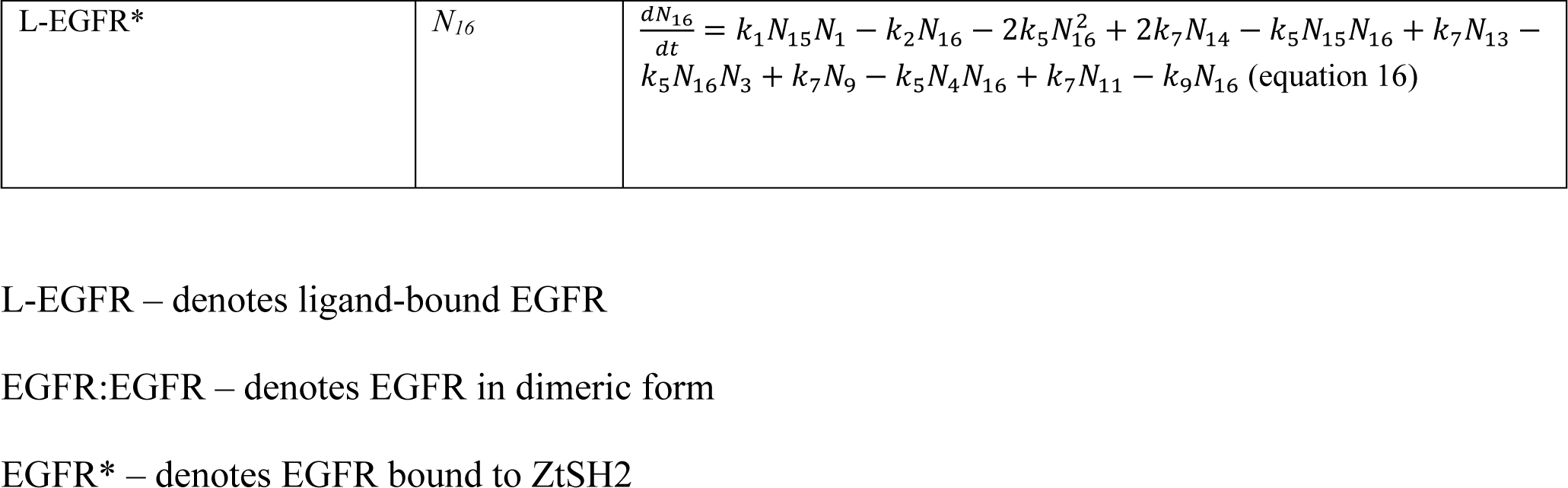

### Parameters

We then parameterized our model using values reported previously. Rate constants and/or binding affinities for every step of the model were found, except for the joint dimerization/phosphorylation of receptors, which we estimated to qualitatively match the kinetics of pYtag responses observed in our experiments.

| Parameter | Notation | Value | Units | Notes |
| --- | --- | --- | --- | --- |
| Receptor-ligand binding | $k_1$ | 0.03 | $\text{nM}^{-1}\text{s}^{-1}$ | (Schoeberl et al., 2002) |
| Ligand dissociating from L-EGFR ($\beta = 1$ corresponds to EGF) | $k_2$ | $\beta * 6.6\text{e-}3$ | $\text{s}^{-1}$ | (Macdonald and Pike, 2008) |
| Ligand dissociating from L-EGFR:EGFR ($\beta = 1$ corresponds to EGF) | $k_3$ | $\beta * 5.7\text{e-}3$ | $\text{s}^{-1}$ | (Macdonald and Pike, 2008) |
| Ligand dissociating from L-EGFR:L-EGFR | $k_4$ | 0.087 | $\text{s}^{-1}$ | (Macdonald and Pike, 2008) |
| Receptor dimerization | $k_5$ | $1\text{e-}5$ | $\text{nM}^{-1}\text{s}^{-1}$ | Estimated |
| EGFR:EGFR dissociation | $k_6$ | $\alpha * 1\text{e-}4$ | $\text{s}^{-1}$ | Estimated |
| L-EGFR:EGFR dissociation ($\gamma = 1$ corresponds to EGF) | $k_7$ | $\gamma * 1\text{e-}4$ | $\text{s}^{-1}$ | Estimated |
| ZtSH2 binding to receptor | $k_8$ | 5 | $\text{nM}^{-1}\text{s}^{-1}$ | Assumed to be large enough that ZtSH2 binding is not rate- limiting, given experimental results. |
| ZtSH2 dissociating from receptor | $k_9$ | 16.67 | $s^{-1}$ | (Ottinger et al., 1998) |
| Scaling parameter for ligand-free dimerization | $\alpha$ | Varies | Unitless | |
| Scaling parameter for ligand-receptor binding | $\beta$ | Varies | Unitless | |
| Scaling parameter for dimerization of ligand-bound receptors | $\gamma$ | Varies | Unitless | |

### Initial conditions

Initial concentrations of EGFR-containing species (*N_3_*-*N_16_*) were determined for the case in which soluble ligand (*N_1_*) is absent and no ZtSH2 is bound to receptors. In this case, the concentrations of ligand-bound and ZtSH2-bound EGFR (*N_4_*; *N_6_*-*N_16_*) were set to zero. The concentration of total EGFR (*N_E,o_*) was calculated by dividing the number of EGFR molecules per cell (assumed to be 250,000 cell^-1^) (Herbst, 2004) by the occupied volume of the cell (treated as a sphere of 10 µm radius). Total EGFR and ZtSH2 were assumed to exist at a 1:1 molar ratio. In the absence of ligand, our model requires that EGFR is confined to the following two states: as a monomer (*N_3_*) or as a ligand-free dimer (*N_5_*). The initial concentrations of *N_3_* and *N_5_* were determined analytically. *N_E,o_* is the sum of the initial concentrations of monomeric EGFR (*N_3,o_*) and EGFR in ligand-free dimers (*N_5,o_*):

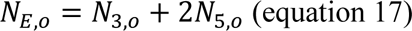

Setting the concentrations of all EGFR-containing species except for *N_3,o_* and *N_5,o_* to zero, equations 3, 5, and 17 yield the following relationship between monomeric EGFR and its ligand-free dimer:

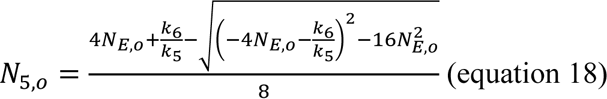

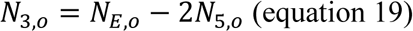

### Altering scaling parameters in model

#### Tuning ligand-free dimerization affinity through α

To assess the effect of ligand-free dimers, α was varied over 2 orders of magnitude; for each value of α, pYtag responses to 20 ng/mL EGF were simulated. Based on these simulations, α was set to 50, since this condition was in qualitative agreement with the biphasic pYtag response observed in experiments.

#### Tuning properties of EGFR ligands through β and γ

To systematically test the effects of ligand-receptor binding and the dimerization affinity of ligand-bound receptors, β and γ were increased by 1000- and 100-fold respectively, and pYtag responses were simulated for ligand doses ranging from 0-5000 ng/mL.

#### Simulating effects of GBM mutants

The responses of WT EGFR and GBM-associated mutants were predicted in response to 20 ng/mL EREG. As described above, ligand-receptor binding affinity and ligand-bound dimerization affinity were decreased by increasing β and γ, respectively. To simulate WT EGFR exposed to EREG, we increased β 50-fold and increased γ 100-fold, relative to simulations of EGF treatment at the same ng/mL dose of ligand; the simulated response under these conditions was in qualitative agreement with the rapid, transient peak and transient plateau of signaling observed in experiments (**Figures 3A and 3E**). To simulate GBM-associated mutants treated with EREG, we increased β 6-fold and increased γ 650-fold (Hu et al., 2022), relative to WT EGFR treated with EREG.

## Quantification and statistical analysis

### Subtraction of background fluorescence from images

All images displayed in figures, and images used in subsequent analyses, were subjected to subtractions of background fluorescence either using a flat-field correction or by subtracting the intensity of cell-devoid regions from raw TIFF files. Images except for those of MCF10A cells on synthetic substrata were subjected to flat-field corrections. To perform a flat-field correction, raw TIFF files were imported into FIJI and subtracted of background fluorescence using a gaussian-blurred image of a sample containing cell culture medium but lacking cells. For images of MCF10A cells on synthetic substrata, the mean gray value (intensity) of a region absent of cells was measured and subtracted from each pixel of the image at the same time point.

### Frame averaging for images presented in Figure 6D

For the mScarlet-ZtSH2 images presented in Figure 6D, each image shows the average of two successive frames to reduce background noise. The full movie without averaging is also included as Video S5.

### Quantification of pYtag and Grb2 biosensor responses

Cytosolic regions of randomly selected cells positive for both the fluorescently-labeled RTK(s) and pYtag/Grb2 reporter(s) of interest were segmented in FIJI and the mean intensity was measured at each time point. In rare cases, abnormally dark or bright images were captured by the confocal microscope, causing sudden spikes in the cytosolic intensities measured in all cells; measurements from these aberrant images were rejected, and the cytosolic intensities of cells at the previous time point were used as a placeholder. CSV files containing cytosolic intensities of pYtag/Grb2 reporters were exported to R for subsequent analysis.

Raw cytosolic intensities of pYtag/Grb2 reporters were normalized to quantify the percentage of reporter cleared from the cytosol after stimulation with ligand. After validating each pYtag biosensor, the percentage of reporter cleared from the cytosol is hence referred to as the activity of the RTK of interest, *activity*_*RTK*_(*t*):

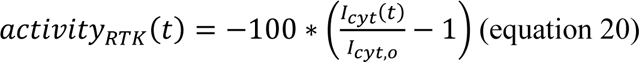

where *I*_*cyt*_(*t*) is the cytosolic intensity of the reporter at a given time point, and *I*_cyt,o_ is the mean cytosolic intensity of the reporter prior to stimulation with ligand. Mean pYtag/Grb2 responses over time were then calculated as the mean ± S.D. of population-averaged means from each experiment.

### Quantification of ZtSH2 and EGFR membrane localization in MCF10A cells

Enrichment of ZtSH2 and EGFR at cell membranes in Figure 2 was quantified in FIJI. The Straight Line feature was used to draw a region of interest intersecting perpendicularly with either media-exposed membranes or cell-cell contacts. Intensity profiles for ZtSH2 and EGFR channels were then measured using the Plot Profile feature and exported to R for analysis.

### Quantification of pYtag and ErkKTR responses in MCF10A cells

Individual *z*-slices from 3D timelapse imaging were used to quantify both EGFR pYtag and ErkKTR responses in MCF10A cells cultured on soft substrata. EGFR pYtag responses were quantified as described above (see **Quantification of pYtag and Grb2 biosensor responses**).

Using the ErkKTR channel, nuclear and cytoplasmic regions of individual cells were segmented in FIJI and the intensity of these regions was measured at each time point. ErkKTR-reported Erk activity was calculated by dividing the cytosolic KTR intensity by the nuclear KTR intensity at each time point. Each cell’s EGFR pYtag and ErkKTR trajectories were normalized to their respective minimum and maximum readouts for the reporter of interest.

### Statistical analysis and replicates

All experiments were performed over at least 3 biological replicates or 2 independent experiments. Biological replicates are defined as biologically distinct samples aimed to capture biological variation. Independent experiments are defined as biologically distinct samples prepared and analyzed on separate days.

## Supplementary Figures

**Figure S1.**
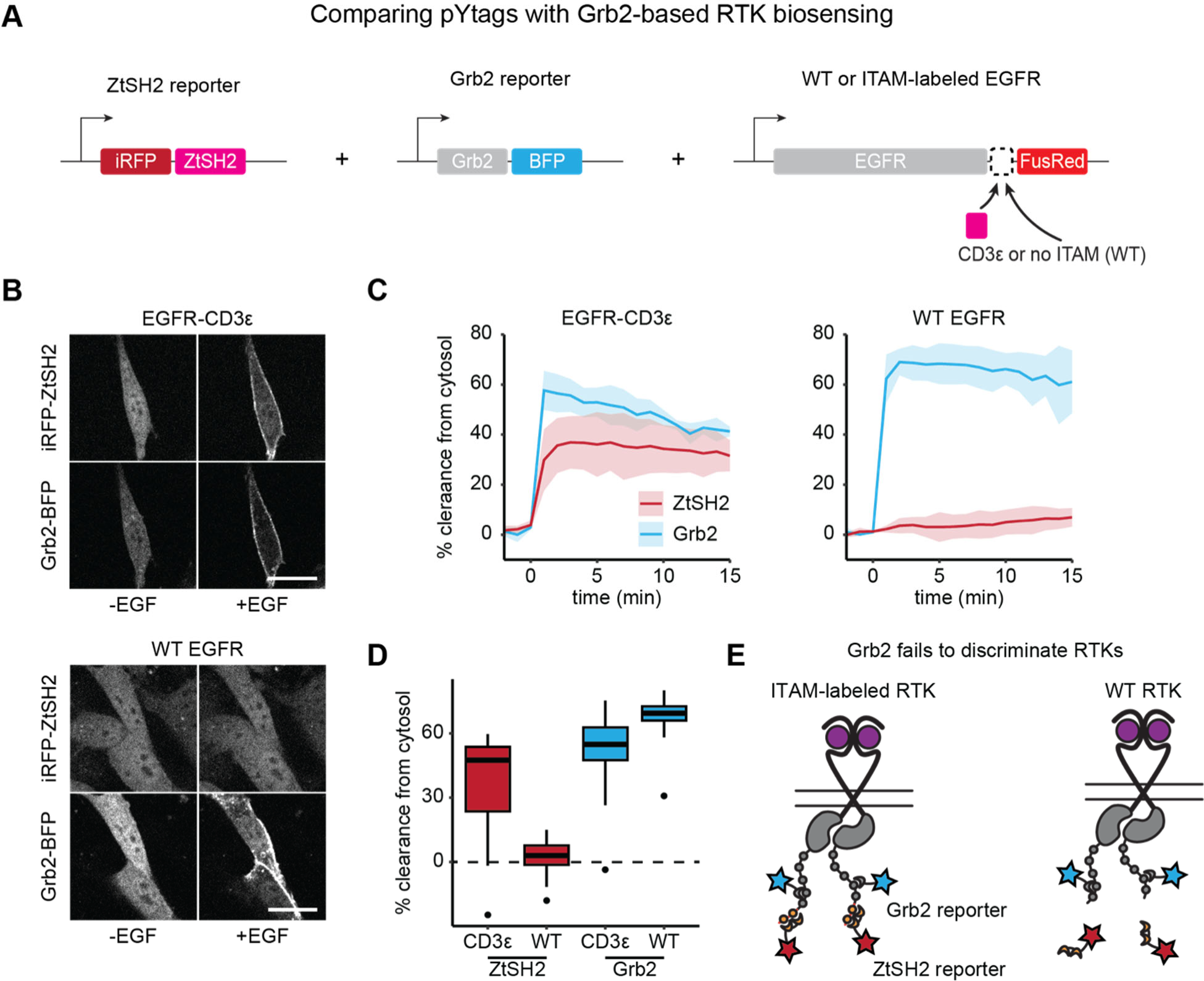
Grb2 fails to discriminate between ITAM-labeled and unlabeled RTKs. (**A**) ZtSH2- and Grb2-based reporters of RTK signaling were co-expressed in cells expressing either ITAM-labeled or WT EGFR. (**B**) NIH 3T3 cells before and 5 min after treatment with EGF (100 ng/mL). Scale bars, 20 µm. (**C**) Mean ± S.D. clearance of ZtSH2 and Grb2 reporters from the cytosol following treatment with EGF (100 ng/mL). *n* = 3 independent experiments. (**D**) Clearance of reporters from the cytosol 5 min after treatment with EGF in (C). Lines denote mean values, boxes denote 25-75^th^ percentiles, and whiskers denote minima and maxima. For each condition, *n* > 16 cells from 3 independent experiments. (**E**) ZtSH2 reports the signaling of ITAM-labeled RTKs, while Grb2 fails to discriminate between RTKs displaying or lacking ITAMs.

**Figure S2.**
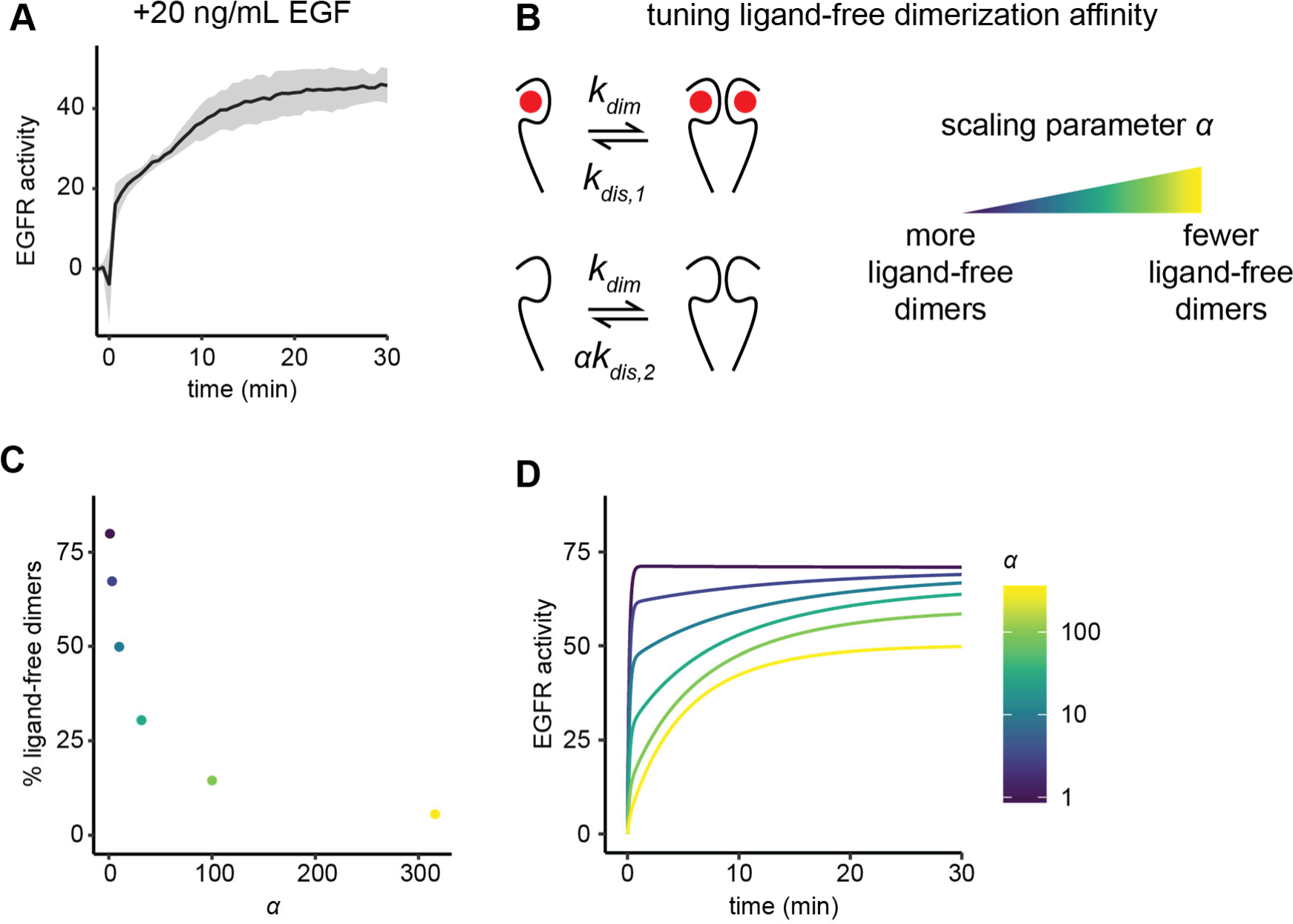
Ligand-free dimers in mathematical model recapitulate biphasic signaling response of EGFR. (**A**) EGFR pYtag response to EGF (20 ng/mL) shown in Figure 2B. (**B**) The effect of ligand-free dimers on EGFR pYtag responses was simulated by tuning the dissociation rate of ligand-free dimers using scaling parameter α. (**C**) Percentage of receptors existing as ligand-free dimers before ligand stimulation as a function of α. Color of data points corresponds to color of curves in (D). (**D**) Simulated EGFR pYtag responses to EGF for varying values of α.

**Figure S3.**
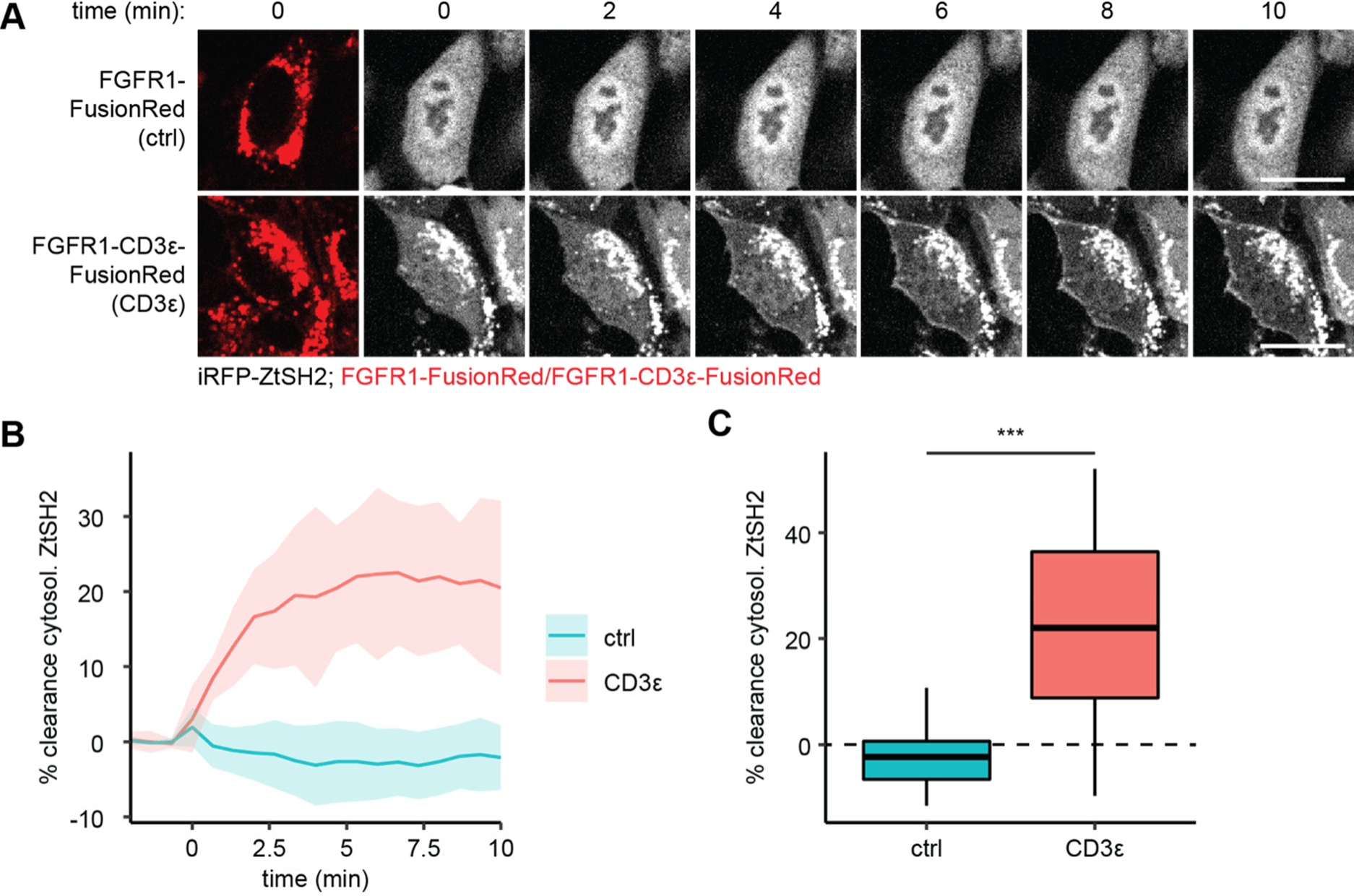
A pYtag-based biosensor of FGFR1 signaling. (**A**) Images of NIH 3T3 cells co-expressing iRFP-ZtSH2 with either FGFR1-FusionRed or FGFR1-CD3ε-FusionRed, treated with FGF4 (100 ng/mL). Scale bars, 20 µm. (**B**) Mean ± S.D. clearance of cytosolic ZtSH2 in cells expressing FGFR1-FusionRed (ctrl) or FGFR1-CD3ε-FusionRed (CD3ε) from (A). *n* = 3 independent experiments. (**C**) Clearance of ZtSH2 from the cytosol 10 min after treatment with FGF4 in (B). Lines denote mean values, boxes denote 25-75^th^ percentiles, and whiskers denote minima and maxima. For each condition, *n* > 38 cells from 3 independent experiments. *** *p* < 0.001 by Kolmogorov-Smirnov test.

**Figure S4.**
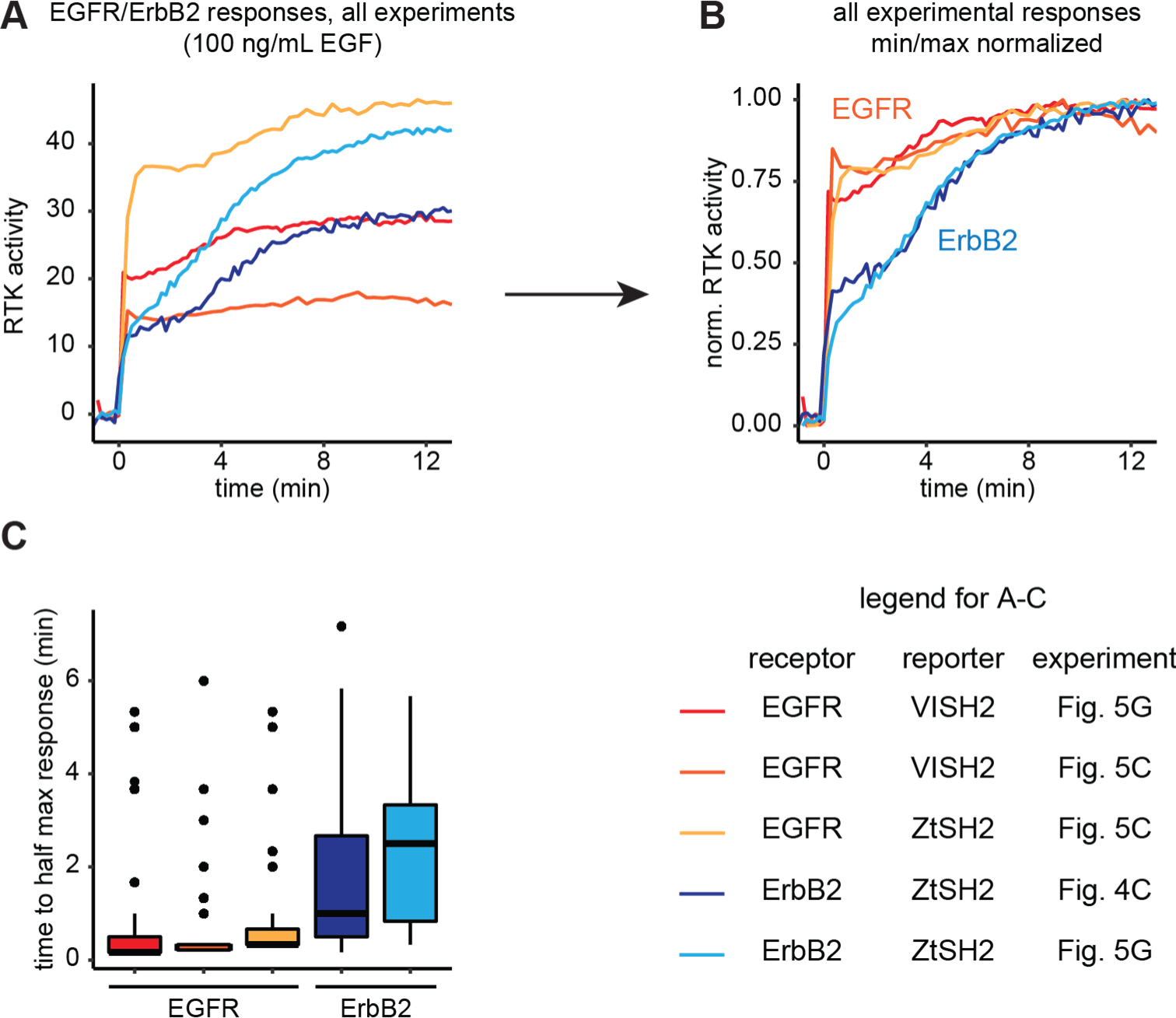
Comparison of EGFR and ErbB2 responses across experiments. (**A**) Mean responses of EGFR and ErbB2 to EGF (100 ng/mL) were collected from experiments with or without multiplexed biosensors (see legend at bottom right of figure) and (**B**) normalized to their minimum and maximum responses for comparison. (**C**) Time to half maximal response from (B). Lines denote mean values, boxes denote 25-75^th^ percentiles, and whiskers denote minima and maxima. For each condition, *n* ≥ 30 cells from 3 independent experiments.

## Supplementary Video legends

**Video S1.** Timelapse of iRFP-ZtSH2 in NIH 3T3 cells co-expressing iRFP-ZtSH2 and EGFR- CD3ε-FusionRed. Cells were first treated with EGF (100 ng/mL) then treated Gefitinib (10 µM) at the times denoted in the video. Related to Figure 1C.

**Video S2.** Max intensity projection timelapse images of MCF10A cells co-expressing EGFR pYtag and ErkKTR, cultured on soft substrata and treated with EGF (100 ng/mL). Related to Figure 2F.

**Video S3.** Timelapse of NIH 3T3 cells co-expressing VISH2 and ZtSH2 reporters, and either SLP76- or CD3ε-labeled EGFR, treated with EGF (100 ng/mL). Related to Figure 5B.

**Video S4.** Timelapse of NIH 3T3 cells co-expressing pYtag biosensors for EGFR and ErbB2, treated with EGF (100 ng/mL). Related to Figure 5F.

**Video S5.** Timelapse of HEK 293T cells expressing mScarlet-ZtSH2 and an endogenously labeled EGFR-CD3ε-mNeonGreen, treated with EGF (100 ng/mL). Left panel shows mScarlet fluorescence; right panel shows mNeonGreen fluorescence at the indicate time points. Scale bar, 20 μm. Related to Figure 6D.

